# Presynaptic cAMP-PKA-mediated potentiation induces reconfiguration of synaptic vesicle pools and channel-vesicle coupling at hippocampal mossy fiber boutons

**DOI:** 10.1101/2023.12.04.569907

**Authors:** Olena Kim, Yuji Okamoto, Nils Brose, Ryuichi Shigemoto, Peter Jonas

## Abstract

It is widely believed that information storage in neuronal circuits involves nanoscopic structural changes at synapses, resulting in the formation of synaptic engrams. However, direct evidence for this hypothesis is lacking. To test this conjecture, we combined chemical potentiation, functional analysis by paired pre-postsynaptic recordings, and structural analysis by electron microscopy (EM) and freeze-fracture replica labeling (FRL) at the hippocampal mossy fiber synapse, a key synapse in the trisynaptic circuit of the hippocampus. Biophysical analysis of synaptic transmission revealed that forskolin-induced chemical potentiation increased the readily releasable vesicle pool size (RRP) and vesicular release probability (P_r_) by 146% and 49%, respectively. Structural analysis of mossy fiber synapses by EM and FRL demonstrated an increase in the number of vesicles close to the plasma membrane and the number of clusters of the priming protein Munc13-1, indicating an increase in the number of both docked and primed vesicles. Furthermore, FRL analysis revealed a significant reduction of the nearest neighbor distance (NND) between Munc13-1 and Ca_V_2.1 Ca^2+^ channels, suggesting reconfiguration of the channel-vesicle coupling nanotopography. Our results indicate that presynaptic plasticity is associated with structural reorganization of active zones (AZs). We propose that changes in potential nanoscopic organization at synaptic vesicle release sites may be correlates of learning and memory at a plastic central synapse.

## INTRODUCTION

Synapses are key sites of exchange and storage of information in the brain (Abbott and Regehr, 2004). Transmitter release has been extensively characterized by biophysical techniques, going back to classical work at the neuromuscular junction (Katz, 1969) and more recent studies at the calyx of Held in the auditory brainstem (Neher, 2017). In parallel, synaptic structure has been characterized in much detail by light and electron microscopy (EM; Frotscher et al., 2006; Harris, 2020). However, the connection between synaptic biophysics and morphology remains unclear. For example, biophysical analysis often determines the number of functional release sites, but the structural correlates have not been identified (Holler et al., 2021). Furthermore, biophysical measurements report the coupling distance between Ca^2+^ channels and release sensors (Adler et al., 1991; Eggermann et al., 2012), but the physical correlate of this measure remains undefined. Finally, biophysical studies often delineate different vesicle pools at presynaptic active zones (AZs), specialized sites of synaptic vesicles fusion and neurotransmitter release (Rizzoli and Betz, 2005; Alabi and Tsien, 2012; Neher, 2015; Kaeser and Regehr, 2017; Emperador-Melero and Kaeser, 2020), but how these pools correlate to vesicle populations in electron micrographs or tomograms remains unclear. For example, the readily releasable pool (RRP) often tightly correlates with the pool of docked vesicles (i.e. vesicles in the direct contact with AZ plasma membrane; Imig et al., 2014), but exceptions to this correlation have been also observed (Man et al., 2015; Wang et al., 2016; Tan et al., 2022). Further correlated biophysical-structural approaches are needed to address these questions.

Transmitter release at central synapses is not constant but undergoes substantial activity-dependent changes (Monday et al., 2018). Structural changes associated with synaptic plasticity are of particular interest, because they may represent parts of “engrams”, defined as physical, chemical, or structural changes underlying information storage in the brain (Josselyn and Tonegawa, 2020). However, because of the nanoscopic scale of the modifications, the nature of these engrams has long remained elusive (Lashley, 1950). Recent work suggested that presynaptic short-term potentiation associated with an increase in RRP and release probability (P_r_, the probability of synaptic vesicle fusion with the plasma membrane) leads to the formation of “pool engrams” (Vandael et al., 2020). However, changes in Ca^2+^ channel localization or channel–vesicle coupling may occur in parallel and could affect the P_r_ of SVs. Moreover, whether similar changes in RRP occur during more long-lasting forms of plasticity or through the effects of neuromodulators remains to be determined (Patzke et al., 2019). To understand the precise nature of synaptic engrams, a nanoscale analysis of the topographical arrangement of presynaptic Ca^2+^ channels and docked synaptic vesicles (SVs) before and after plasticity induction is needed. Although recent work demonstrated the feasibility of such a technically demanding approach (Aldahabi et al., 2022), rigorous correlated biophysical and morphological analysis at defined glutamatergic synapse at the unitary level is lacking.

The hippocampal mossy fiber–CA3 pyramidal neuron synapse, formed between dentate gyrus granule cells (GCs) and CA3 pyramidal neurons (PNs), is an ideal synapse to tackle these questions (Nicoll and Schmitz, 2005; Bischofberger et al., 2006a). First, it is accessible for direct presynaptic–postsynaptic recording (Vandael et al., 2021), which enables precise biophysical analysis of transmission at the single-synapse level. Second, it has a unique extent of presynaptic plasticity, including facilitation, post-tetanic potentiation (PTP; Vandael et al., 2020), and long-term potentiation (LTP; Salin et al., 1996). Third, the plasticity at hippocampal mossy fiber synapses is dependent on well-defined canonical signaling pathways. For example, high-frequency stimulation (HFS) induces several cyclic adenosine monophosphate (cAMP)-dependent forms of presynaptic plasticity (Huang et al., 1994; Weisskopf et al., 1994; Tong et al., 1996; López-García et al., 1996; Maccaferri et al., 1998; Midorikawa and Sakaba 2017; Vandael et al., 2020). Similarly, forskolin, an adenylyl cyclase (AC) activator, leads to marked chemical potentiation (Weisskopf et al., 1994; Maccaferri et al., 1998; Midorikawa and Sakaba 2017). Finally, it is accessible to quantitative structural analysis by EM (Rollenhagen et al., 2007; Wilke et al., 2013), high-pressure freezing (HPF; Studer et al., 2014; Borges-Merjane et al., 2020; Imig et al., 2020), and freeze-fracture replica labeling (FRL; Hagiwara et al., 2005).

Using combined biophysical and structural analysis, we found that AC activation by forskolin increased the size of the RRP and the docked vesicle pool in parallel. Furthermore, we discovered that forskolin changed the distance between Ca_V_2.1 (P/Q-type) Ca^2+^ channels and Munc13-1, an essential vesicle priming protein and putative marker of primed vesicles (Augustin et al., 1999a, 1999b; Varoqueaux et al., 2002; Varoqueaux et al., 2005; Lipstein et al. 2013; Imig et al., 2014; Sakamoto et al., 2018). Taken together, our results provide novel insights into the mechanisms of chemical potentiation at central synapses and the nanoscopic mechanisms of engram formation.

## RESULTS

### Activation of the cAMP-PKA signaling pathway increases RRP and P_r_

HFS-induced potentiation has been demonstrated to increase the RRP size (Vandael et al., 2020). However, the mechanisms of chemical potentiation remain controversial. Recent work suggested an increase in number of release sites (Orlando et al., 2021) or an accumulation of presynaptic Ca^2+^ channels (Fukaya et al., 2021), but delineating the exact mechanisms requires precise analysis of synaptic transmission at the unitary level.

To pinpoint the biophysical mechanisms of chemical potentiation at hippocampal mossy fiber synapses, we performed paired recordings from presynaptic mossy fiber boutons (MFBs) and postsynaptic CA3 PNs (Figure 1). To maximally activate the cAMP pathway, we applied forskolin, a widely used AC activator (Huang et al., 1994; Weisskopf et al., 1994; Tong et al., 1996; López-García et al., 1996). 5- min application of 50 µM forskolin caused a potentiation of evoked excitatory postsynaptic currents (EPSCs) at MFB–CA3 PN synapses to 368% of control amplitude (Figures 1A–C; EPSC_1_ – control: 154.1 ± 15.1 pA (mean ± SEM), median 146.8 pA; forskolin: 566.9 ± 122.3 pA, median 511.8 pA, n = 8 pairs and N = 8 rats in all analysis; control vs. forskolin: P = 0.0078, Wilcoxon signed-rank test). These results confirm that chemical potentiation by forskolin markedly increases the strength of mossy fiber synaptic transmission at the single-synapse level.

**Figure 1.**
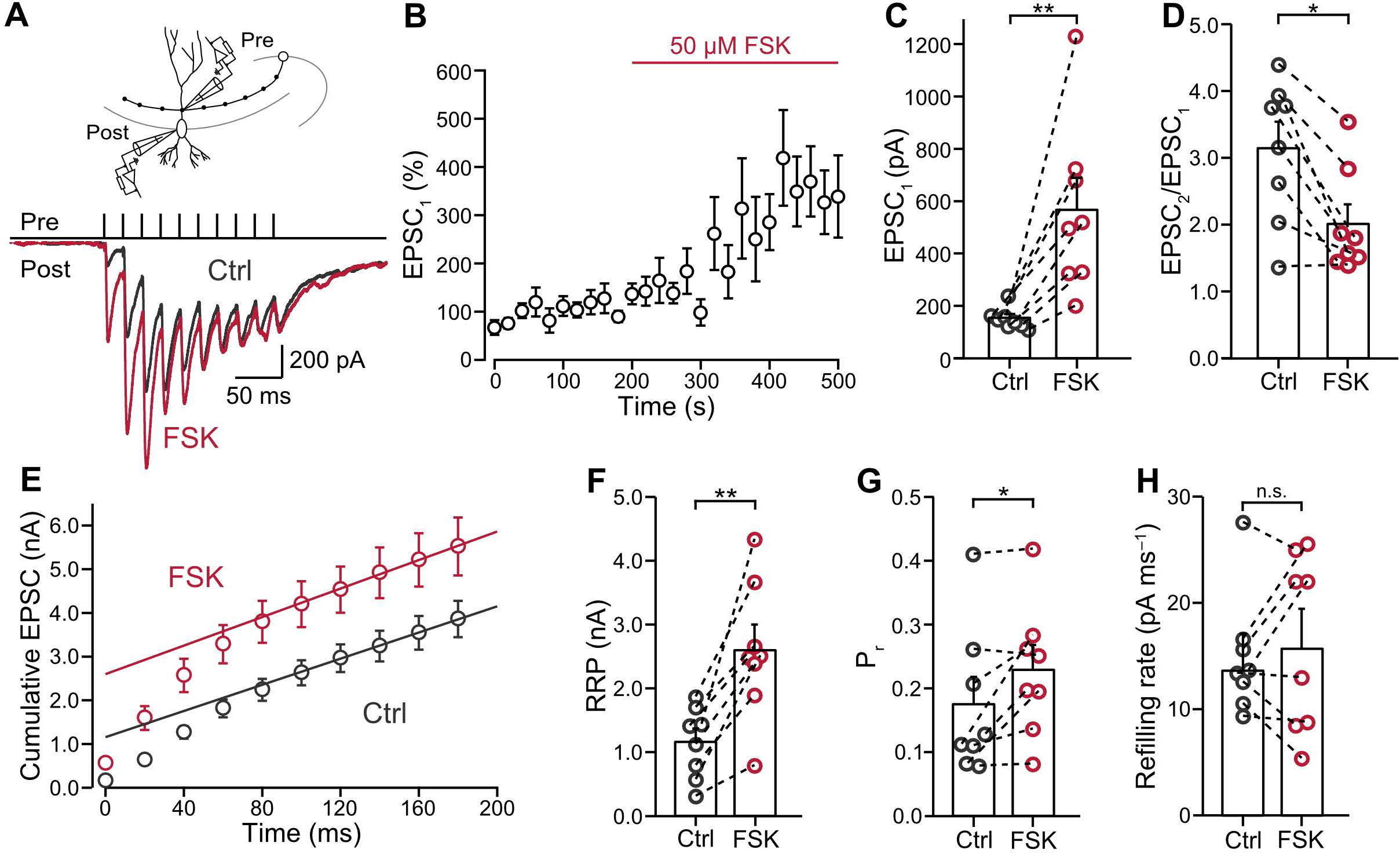
Forskolin-induced chemical potentiation is primarily mediated by an increase in RRP size. (A) Top panel: schematic illustration of the paired recording. Single mossy fiber boutons (MFBs) were stimulated in the tight-seal, cell-attached configuration, while postsynaptic CA3 pyramidal neurons (CA3-PNs) were simultaneously recorded in the whole-cell voltage-clamp configuration. Center panel: 50-Hz train of ten stimuli. Bottom panel: overlay of average excitatory postsynaptic currents (EPSCs) before (“Ctrl”; gray) and in the presence of 50 µM forskolin (“FSK”; red). (B) Normalized EPSC_1_ peak amplitudes plotted against experimental time. Red horizontal line indicates the application time of forskolin (“FSK”). Note that the onset of forskolin application is the time point of switching solutions from forskolin-free artificial cerebrospinal fluid (ACSF) to forskolin-containing ACSF. Forskolin-containing ACSF reached the recording chamber about 60–90 seconds after the onset of forskolin application. Data from 8 pairs (8 rats). (C) Summary bar graph of EPSC_1_ peak amplitudes before (gray) and in the presence of 50 µM forskolin (red). Bars and whiskers show mean + SEM. (D) Summary bar graph of paired-pulse ratio (EPSC_2_ / EPSC_1_) before (gray) and in the presence of 50 µM forskolin (red). Bars and whiskers show mean + SEM. (E) Cumulative plot of EPSC peak amplitudes during a 50-Hz train with 10 stimuli before (“Ctrl”, gray) and in the presence of 50 µM forskolin (“FSK”, red). Data points during the last 4 stimuli (at time points ≥ 120 ms) were fit by linear regression and back-extrapolated to time point 0. **(F–H)** Summary bar graphs of readily releasable pool size (RRP; F), release probability (P_r_; G), and refilling rate (H), estimated from the cumulative EPSC plot (E), before (“Ctrl”, gray) and in the presence of 50 µM forskolin (“FSK”, red). Bars and whiskers show mean + SEM. Note that chemical potentiation induces a large increase in RRP size and a small increase in P_r_. See Figure S1.

To determine the locus of chemical potentiation, we analyzed the paired-pulse ratio (PPR; EPSC_2_/EPSC_1_; Figure 1D). PPR significantly decreased after forskolin application (EPSC_2_/EPSC_1_ – control: 3.14 ± 0.40 (mean ± SEM), median 3.47; forskolin: 2.01 ± 0.29, median 1.70; control vs. forskolin: P = 0.0156, Wilcoxon signed-rank test), consistent with a presynaptic locus of potentiation and an increase in release probability. Furthermore, we analyzed amplitude and frequency of miniature EPSCs (mEPSCs; Figure S1). mEPSCs frequency but not amplitude was affected by forskolin (Figure S1D–G, mEPSCs frequency – control: 3.09 ± 0.61 Hz (mean ± SEM), median 2.07 Hz; forskolin: 4.33 ± 0.54 Hz, median 4.04 Hz; control vs. forskolin: P = 0.0185; mEPSCs amplitude control: 24.7 ± 1.5 pA, median 23.6 pA; forskolin: 23.6 ± 0.9 pA, median 23.1 pA; control vs. forskolin: P = 0.4131, Wilcoxon signed-rank test; n = 11 cells and N = 3 rats). These results suggest a presynaptic locus of forskolin-induced potentiation.

To dissect the biophysical mechanisms of forskolin potentiation, we performed analysis of cumulative EPSC amplitudes (Figure 1E; Schneggenburger et al., 1999; Neher, 2015; Vandael et al., 2020). This method can distinguish between changes in RRP and P_r_, although the resulting RRP values represent “pool decrement” rather than absolute pool size (see Methods; Neher, 2015). Notably, we found that both RRP and P_r_ significantly increased after forskolin application (Figures 1F and G; RRP – control: 1.16 ± 0.21 nA (mean ± SEM), median 1.27 nA; forskolin: 2.60 ± 0.40 nA, median 2.50 nA; control vs. forskolin: P = 0.0078, Wilcoxon signed-rank test; P_r_ control: 0.17 ± 0.04 (mean ± SEM), median 0.12; forskolin: 0.23 ± 0.04, median 0.22; control vs. forskolin: P = 0.0391, Wilcoxon signed-rank test). In contrast, the refilling rate remained unchanged after forskolin application (Figure 1H; refilling rate – control: 15.0 ± 2.1 pA ms^−1^ (mean ± SEM), median 13.6 pA ms^−1^; forskolin: 16.3 ± 3.1 pA ms^−1^, median 17.5 pA ms^−1^; control vs. forskolin: P = 0.7422, Wilcoxon signed-rank test). Thus, chemical potentiation primarily involved an increase in pool size, but also an increase in P_r_, reminiscent of hippocampal mossy fiber PTP (Vandael et al., 2020).

To further assess the functional significance of chemical potentiation, we tested the effects of forskolin on unitary excitatory postsynaptic potentials (EPSPs; Figure 2A). Forskolin increased the amplitude of EPSP_1_ (Figure 2B; EPSP_1_ – control: 4.39 ± 1.99 mV (mean ± SEM), median 1.35 mV; forskolin: 9.56 ± 2.42 mV, median 11.88 mV, n = 7 pairs and N = 7 rats; control vs. forskolin: P = 0.0313, Wilcoxon signed-rank test). However, the amplitude of the EPSP remained below the threshold for action potential (AP) initiation in both conditions. In contrast, for the second and third EPSP in a 50-Hz train, the AP probability was markedly increased after forskolin application (Figure 2C; probability of third AP – control: 0.36 ± 0.17 (mean ± SEM); forskolin: 0.69 ± 0.14, n = 7 pairs and N = 7 rats; control vs. forskolin: P = 0.0313, Wilcoxon signed-rank test). Thus, chemical potentiation markedly regulates the conditional detonation properties of the synapse (Henze et al., 2002; Vyleta et al., 2016).

**Figure 2.**
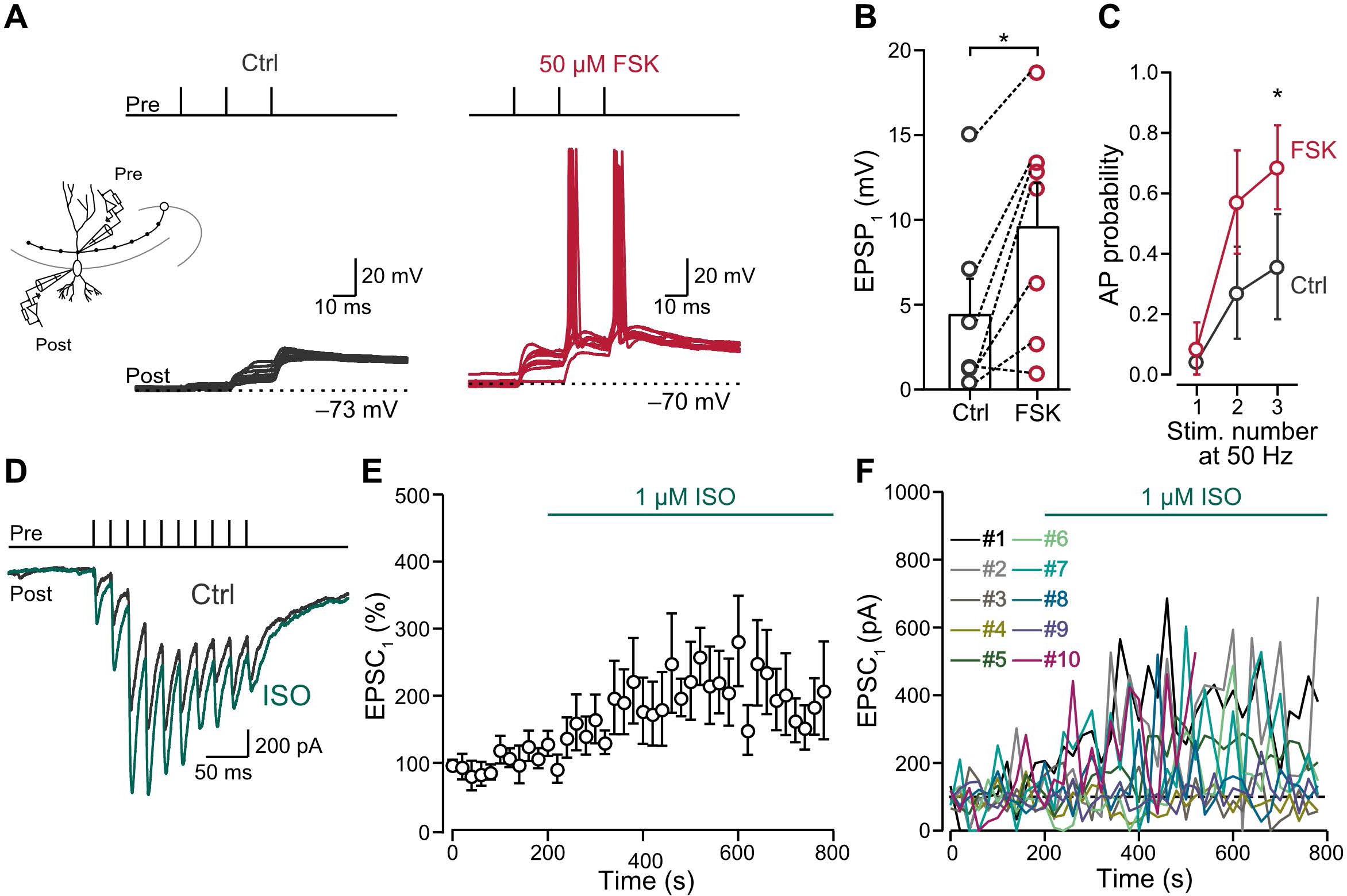
Physiological effects of cAMP-PKA activation in MFB. (A) Side panel: schematic illustration of the paired recording. Single MFBs were stimulated in the tight-seal, cell-attached configuration, while postsynaptic CA3-PNs were simultaneously recorded in the whole-cell current-clamp configuration. Top panel: 50-Hz train of three stimuli. Bottom panel: representative traces of excitatory postsynaptic potentials (EPSPs) and action potentials (APs) in CA3-PNs before (gray) and after 50 μM forskolin (red). (B) Summary bar graph of EPSP_1_ peak amplitude before (“Ctrl”, gray) and in the presence of 50 μM forskolin (“FSK”, red). Bars and whiskers show mean + SEM. (C) Probability of AP firing before (“Ctrl”, gray) and in the presence of 50 μM forskolin (“FSK”, red). (D) Top panel: 50-Hz train of ten stimuli. Bottom panel: representative average EPSCs before (“Ctrl”; gray) and in the presence of 1 µM isoproterenol (“Iso”; green). The recording configuration similar to figure 1A. (E) Normalized EPSC_1_ peak amplitudes plotted against experimental time. Green horizontal line indicates the application time of isoproterenol (“Iso”). The switch of the solution is similar to figure 1B. Data from 10 pairs in 8 rats. (F) Normalized EPSC_1_ peak amplitudes from individual pairs plotted against experimental time. Note that some pairs strongly respond to isoproterenol, whereas others show no effects. Thus, the MFB synapse population is heterogeneous in the sensitivity to the agonist. See Figure S2.

To test whether a more natural activation of the cAMP-PKA signaling pathway had similar effects, we applied 1 µM isoproterenol, a β_1_ and β_2_ adrenergic receptor agonist, known to activate AC via G_s_ proteins (O’Dell et al., 2015; Hansen and Manahan-Vaughan, 2015; Kobayashi et al., 2022; Figures 2D–F and S2). Similar to forskolin, isoproterenol led to potentiation of EPSC_1_ (Figures 2D–E and 2SA; EPSC_1_ – control: 178.9 ± 37.5 pA (mean ± SEM), median 181.4 pA; isoproterenol: 323.6 ± 86.7 pA, median 195.2 pA, n = 10 pairs and N = 8 rats; control vs. isoproterenol: P = 0.0273, Wilcoxon signed-rank test). However, the effects were more variable than those of forskolin, implying heterogeneity in the signaling pathway upstream of the AC (Figure 2F). To further test for the presynaptic locus of this chemical potentiation induced by isoproterenol, we analyzed the PPR (Figure S2B). PPR significantly decreased after isoproterenol application (EPSC_2_/EPSC_1_ – control: 2.97 ± 0.31 (mean ± SEM), median 3.06; isoproterenol: 1.86 ± 0.17, median 1.72; control vs. isoproterenol: P= 0.0009, Wilcoxon signed-rank test), again consistent with a presynaptic locus of potentiation. Further cumulative EPSC analysis revealed that the potentiation led to an increase in RRP size (Figures S2C and D; RRP – control: 1.17 ± 0.26 nA (mean ± SEM), median 1.16 nA; isoproterenol: 1.90 ± 0.38 nA, median 1.46 nA; control vs. isoproterenol: P = 0.0039, Wilcoxon signed-rank test). However, neither P_r_ nor refilling rate changed after induction of chemical potentiation with isoproterenol (Figures S2E and F; P_r_ – control: 0.16 ± 0.02 (mean ± SEM), median 0.15; isoproterenol: 0.17 ± 0.02, median 0.16; control vs. isoproterenol: P = 0.9219, Wilcoxon signed-rank test; refilling rate – control: 12.03 ± 2.26 pA ms^−1^ (mean ± SEM), median 11.13 pA ms^−1^; isoproterenol: 9.65 ± 1.84 pA ms^−1^, median 7.87 pA ms^−1^; control vs. isoproterenol: P = 0.1602, Wilcoxon signed-rank test). These results indicate that natural activation of the cAMP-PKA signaling pathway leads to expansion of the RRP, although with larger variability than direct AC activation.

### Chemical potentiation leads to an expansion of the docked vesicle pool

Considering the robust effects of forskolin on RRP and P_r_, we next wanted to test whether any structural changes occur in the AZ at the vesicle docking level at MFB– CA3 PN synapses (Figure 3). We therefore cryo-fixed acute hippocampal slices with and without forskolin treatment (50 µM; 5 min) and performed freeze-substitution (Figures 3A–D). Forskolin was used in this set of experiments, because of the reliability of the effects and the previously documented compatibility with EM experiments (Patzke et al., 2019; Rey et al., 2020; Orlando et al., 2021). Vesicles, whose outer membrane and presynaptic AZ membrane were in direct contact, were considered “docked”. After forskolin application, the number of docked SVs increased in comparison to control conditions (Figure 3E; number of docked vesicles per 100 nm AZ profile length – control: 0.78 ± 0.52 (mean ± SD), median 0.82, n = 159 AZ, N = 3 mice; forskolin: 1.20 ± 0.60, median 1.17, n = 149 AZ, N = 3 mice; control vs. forskolin: P < 0.0001, Mann-Whitney test). This change was also observed in cumulative distributions, where the forskolin sample group was shifted towards the right (Figure 3F; control vs. forskolin: P < 0.0001, Mann-Whitney test). In contrast, the diameter of docked vesicles did not change upon application of forskolin (Figures 3G–H; docked vesicle diameter – control: 38.1 ± 17.9 nm (mean ± SD), median 32.0 nm, n = 159 AZ, n = 280 vesicles, N = 3 mice; forskolin: 34.9 ± 12.2 nm, median 32.0 nm, n = 149 AZ, n = 382 vesicles, N = 3 mice; control vs. forskolin: P = 0.1689, Mann-Whitney test; control vs. forskolin: P = 0.2259, Mann-Whitney test). The effects of forskolin in the docked vesicle pool were similar to those observed during HFS-induced mossy fiber PTP (Vandael et al., 2020). However, the effects on vesicle diameter were distinct, because docked vesicle diameter was constant in the case of forskolin potentiation, but slightly increased in the case of HFS-induced PTP (Vandael et al., 2020), Thus, direct activation of the cAMP-PKA signaling pathway affects the docking of smaller and larger SVs to the same extent.

**Figure 3.**
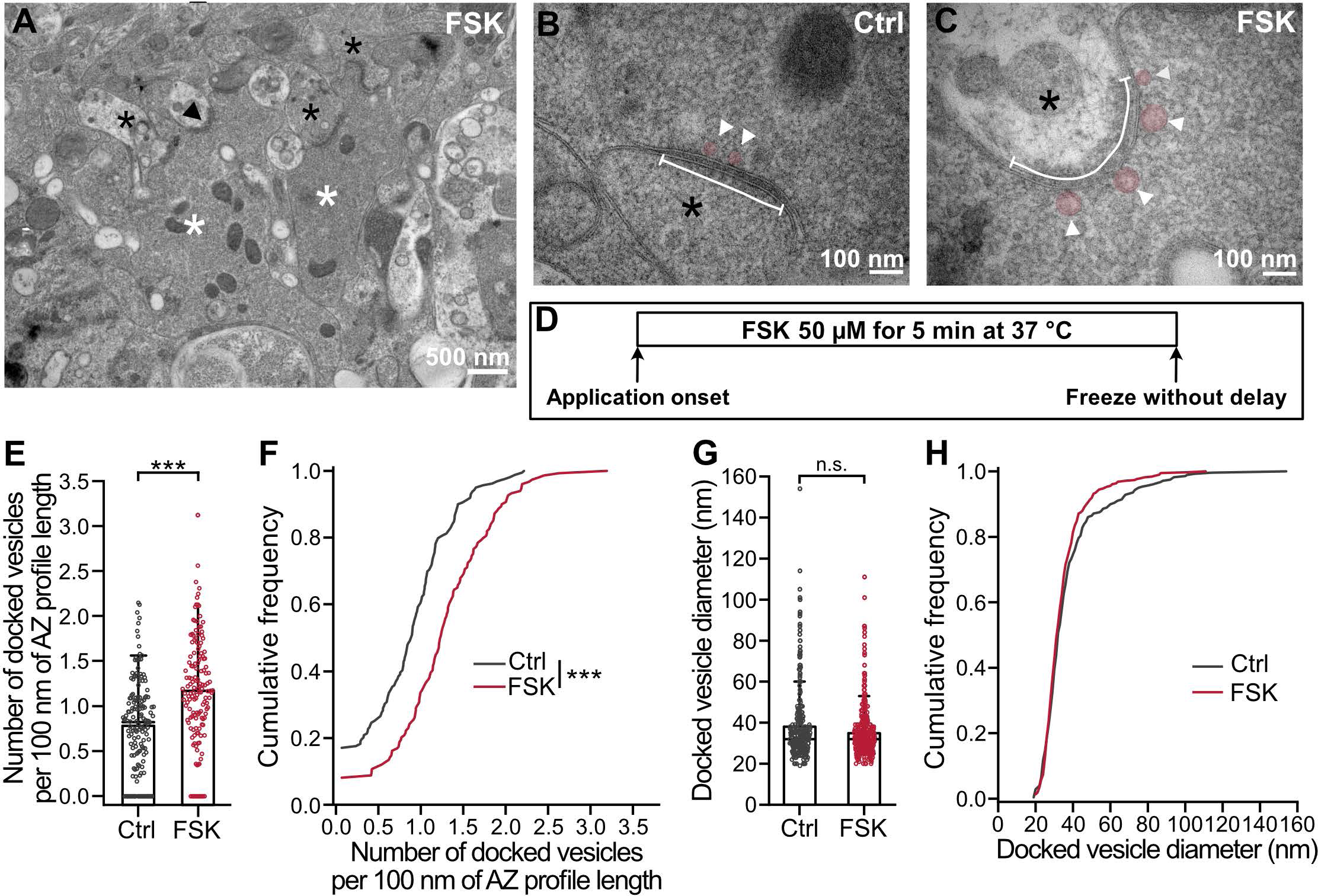
**Docked vesicle pool of MFBs expands after induction of chemical potentiation.** **(A)** Example TEM micrographs from acute hippocampal slices, showing putative MFBs (white asterisks) with putative postsynaptic spines (black asterisks) in *stratum lucidum* of CA3 region. MFBs were recognized based on characteristic morphological features: large bouton size, high density of clear synaptic vesicles (SVs), presence of large dense-core vesicles, high-density of mitochondria, and multiple synaptic contacts with large spines. Black arrowhead points to an AZ shown in (C). **(B** and **C)** Higher magnification TEM micrographs showing AZs (white line) and docked SVs (pink circles and white arrowheads) in putative MFBs in DMSO control (B, “Ctrl”) and after forskolin application (C, “FSK”). **(D)** Schematic representation of the time course of the experiment with 5-min forskolin (FSK) treatment. **(E)** Summary bar graph of the number of docked vesicles per 100 nm of AZ profile length in DMSO control (“Ctrl”, gray) and after forskolin application (“FSK”, red). Bars and whiskers show mean + SD. Horizontal black lines indicate median values. **(F)** Cumulative plots of the data displayed in (E), color scheme is identical to (E). **(G)** Summary bar graph of the diameter of docked vesicles measured in DMSO control (“Ctrl”, gray) and after forskolin treatment (“FSK”, red). Bars and whiskers show mean + SD. Horizontal black lines indicate median values. **(H)** Cumulative plots of the data displayed in (G), color scheme is identical to (G). Scale bar sizes are indicated on the figure panels.

### Chemical potentiation alters the topographical relations between Ca2.1 and Munc13-1

Forskolin may alter the molecular nanoarchitecture of MFB AZs, such as the distribution of Ca^2+^ channels or other AZ proteins in hippocampal MFB terminals (Pauli et al., 2021; Fukaya et al., 2021; Fukaya et al., 2023). To localize membrane-bound proteins with nanometer precision within the AZ, we employed FRL (Figure 4). To probe the Ca^2+^ channel–sensor coupling topography, we targeted Ca2.1, the major type of presynaptic Ca^2+^ channel in MFBs (Castillo et al., 1994; Li et al., 2007), and Munc13-1/2, putative markers of primed vesicles (Varoqueaux et al., 2002; Varoqueaux et al., 2005; Chen et al., 2013; Lipstein et al. 2013; Imig et al., 2014; Sakamoto et al., 2018; Grushin et al., 2022). Although Munc13s do not have transmembrane domains, Munc13-1 in synapses appears to be tightly anchored to the AZ via scaffold proteins (Betz et al. 2001; Limbach et al., 2008) and interactions with components of the sub-membranous cytoskeleton (Sakaguchi et al., 1998), rendering it effectively insoluble (Brose et al., 1995). The corresponding AZ membrane association can be detected by FRL (Rebola et al. 2019; Karlocai et al., 2021; Aldahabi et al., 2022; Grushin et al., 2022). Thus, after cryo-fixation, samples were fractured, and replicas were labeled with anti-Ca2.1 and anti-Munc13-1 antibodies (Figures 4 and S3–S6, Table S1). The density of the labeling of both type of particles remained the same in the presence and absence of forskolin (Figure S4A–C and Table S1; number of particles per 0.1 µm^2^ of AZ area: Ca2.1: control vs. forskolin: P = 0.3598; Munc13-1: control vs. forskolin: P = 0.0689, Mann-Whitney test).

**Figure 4.**
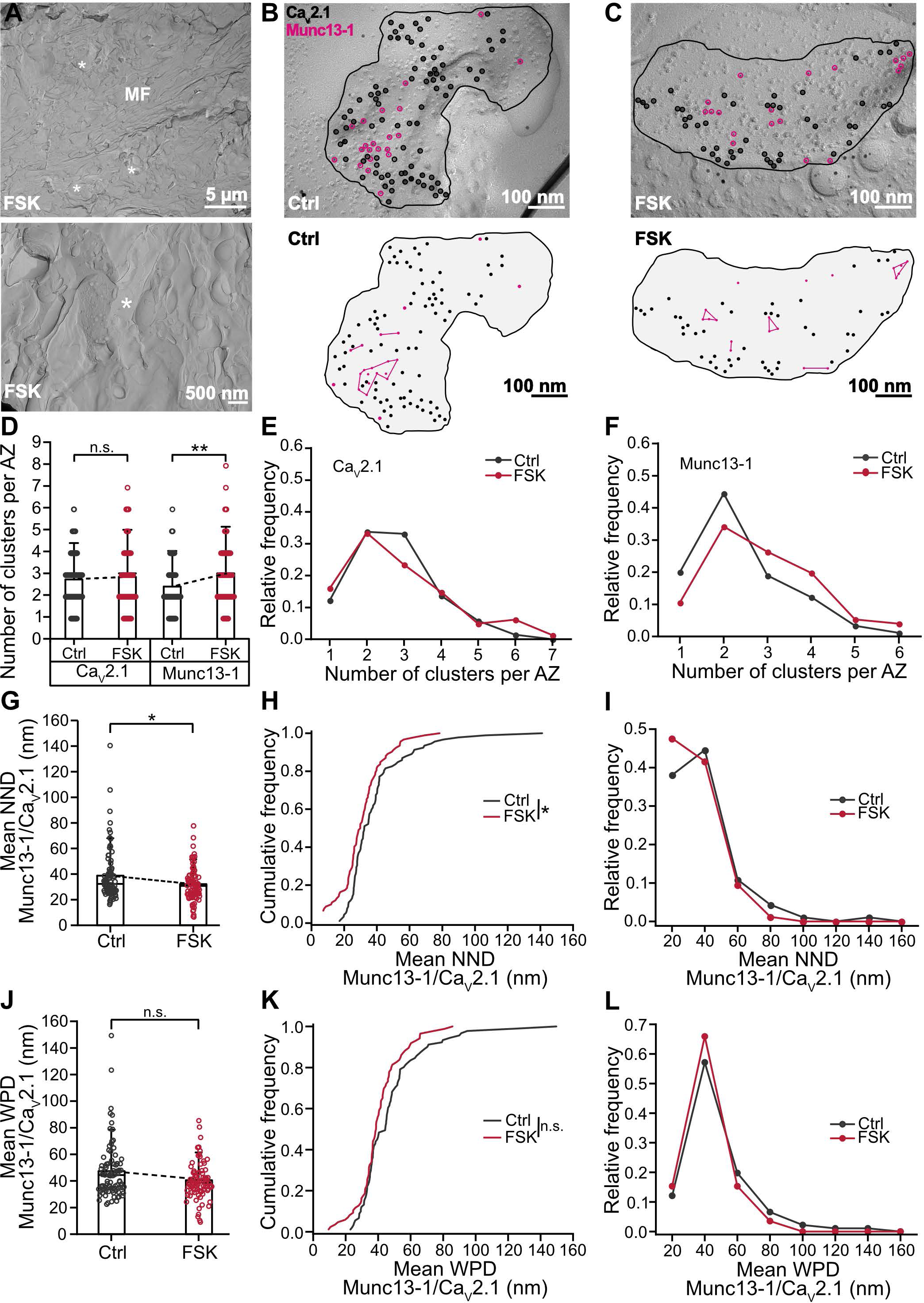
**Rearrangement of Ca_V_2.1 and Munc13-1 proteins within MFB AZs after induction of chemical potentiation.** **(A)** Top panel: example TEM micrographs of freeze-fractured replicas of acute slices showing ultrastructural quality of *stratum lucidum* of the CA3 region, with mossy fiber axon (MF) and putative MFBs (white asterisks) after forskolin (“FSK”). Bottom: Higher magnification TEM micrographs showing putative MFB (white asterisk) in *stratum lucidum* of CA3 region after forskolin (“FSK”). MFBs were recognized based on characteristic morphological features: large bouton size and presence of adjacent cross-fracture with high density of synaptic vesicles (SVs), **(B** and **C)** Top panel: higher magnification micrographs with putative MFB AZs (black line) co-labeled against Ca_V_2.1 (black empty circles) and Munc13-1 (pink empty circles) in DMSO control (B, “Ctrl”) and after forskolin treatment (C, “FSK”). Bottom panel: schematic of the above AZs with Munc13-1 clusters (pink circles connected with pink line) in DMSO control (B, “Ctrl”) and after forskolin treatment (C, “FSK”). Note that some Munc13-1 particles do not belong to any cluster and are considered “noise” points (single pink circles). **(D)** Summary bar graph of the number of Ca_V_2.1 and Munc13-1 clusters per AZ in DMSO control (“Ctrl”, gray) and after forskolin (“FSK”, red). Bars and whiskers show mean + SD. Horizontal black lines indicate median values. **(E–F)** Relative frequency distribution of data displayed in (D), Ca_V_2.1 (E), Munc13-1 (F), color scheme is identical to (D). (G) Summary bar graph of the mean NND between Munc13-1 and Ca_V_2.1 in DMSO control (“Ctrl”, gray) and after forskolin (“FSK”, red). Bars and whiskers show mean + SD. Horizontal black lines indicate median values. (H) Cumulative plots of mean NND between experimental Munc13-1 and Ca_V_2.1 point patterns, color scheme is identical to (G). (I) Relative frequency distribution of data shown in (G), color scheme is identical to (G). (J) Summary bar graph of the mean WPDs between Munc13-1 and Ca_V_2.1, color scheme is identical to (G). Bars and whiskers show mean + SD. Horizontal black lines indicate median values. (K) Cumulative plots of mean WPDs between experimental Munc13-1 and Ca_V_2.1 point patterns, color scheme is identical to (G). (L) Relative frequency distribution of data shown in (J), color scheme is identical to (G). Scale bar sizes are indicated on the figure panels. See Figures S3–S7.

Nearest neighbor distance (NND) and modified Ripley H-function analysis revealed that CaV2.1s and Munc13-1s formed clusters both in the presence and absence of forskolin, as cumulative frequency curves and H-functions of experimental data were significantly different from corresponding curves for randomly distributed particles (Figures S4, S5, and Table S1). DBSCAN analysis (Ester et al., 1996) was used to determine the number of Ca_V_2.1 and Munc13-1 clusters (Figures 4 and S6). The mean number of Ca_V_2.1 clusters did not change in samples cryo-fixed after forskolin treatment (Figures 4D and E and Table S2; control vs. forskolin: P = 0.1595, Mann-Whitney test). In contrast, the number of Munc13-1 clusters increased significantly in samples cryo-fixed after forskolin application (Figures 4D, F and Table S2; control vs. forskolin: P = 0.0088, Mann-Whitney test). Notably, the increase in Munc13-1 clusters agreed with the expansion of docked vesicle pool and RRP after forskolin treatment and HFS (Vandael et al., 2020; Kobbersmed et al., 2020).

The hippocampal MFB synapses are synapses with low initial P_r_ (Jonas et al., 1993; Lawrence et al., 2004; Vyleta et al., 2016). This may be explained by loose coupling between the Ca^2+^ source (Ca^2+^ channels) and the release sensor (synaptotagmin on SVs) in MFBs (Vyleta and Jonas, 2014). To investigate the structural correlates of coupling, FRL experiments with co-labeling of Munc13-1 and Ca_V_2.1 were performed. Interestingly, the mean NND between Munc13-1 and Ca_V_2.1 particles decreased after forskolin treatment (Figures 4G–I and Table S2; control vs. forskolin: P = 0.0275, Mann-Whitney test). Furthermore, the mean weighted pairwise distance (WPD) slightly decreased, although the change was not significant (Figures 4J–L and Table S2; control vs. forskolin: P = 0.0706, Mann-Whitney test). These results suggest that the increase in P_r_ after induction of chemical potentiation could be related to a tightening of the coupling configuration.

After Munc13-1, the second-most abundant isoform expressed at MFB synapses is bMunc13-2 (Augustin et al., 1999a; Varoqueaux et al., 2002; Breustedt et al., 2010). To test the contribution of bMunc13-2 proteins to the priming of SVs during PKA-dependent potentiation, we examined the distribution of bMunc13-2 proteins in MFB AZs (Figures S3–S7 and Table S1). Both NND and Ripley H-function analysis revealed a tendency of bMunc13-2 proteins to cluster within MFB AZs in the presence and absence of forskolin application, albeit significant only in case of the NND (Figure S3–5 and Table S1). Unlike Munc13-1, bMunc13-2 did not show any significant alterations in the cluster number after forskolin application (Figures S7B–C and Table S2; control vs. forskolin: P = 0.8124, Mann-Whitney test). Furthermore, no change in the mean NND and the mean WPD between bMunc13-2 and Ca_V_2.1 was observed (Figures S7D–I and Table S2; NND – control vs. forskolin: P = 0.1231; WPD – control vs. forskolin: P = 0.6687, Mann-Whitney test). These results indicate that the effects of forskolin on localization of Munc13 isoforms were differential, resulting in a selective increase of Munc13-1 but not bMunc13-2 clusters. This is consistent with the hypothesis that the Munc13-1 protein isoform is the major priming factor in hippocampal MFBs, involved in the expression of PKA-dependent potentiation.

### Spatial reorganization of Munc13-1 depends on PKA activity

The effects of forskolin on spatial reorganization of Munc13-1 may be PKA-dependent (Weisskopf et al., 1994), but could also involve other pathways, e.g. Epac (Fernandes et al., 2015). To test whether the changes in Munc13-1 spatial distribution were PKA-dependent, we repeated the experiments in the presence of 10 µM H-89, a PKA inhibitor (Figure 5). In the presence of H-89, the number of Munc13-1 clusters per AZ did not change after forskolin application (Figure 5E and F; Table S3; control vs. H- 89: P = 0.6741, control vs. H-89+forskolin: P = 0.9893, H-89 vs. H-89+forskolin: P = 0.6869, Mann-Whitney test). In addition, the mean NND between Munc13-1 and Ca_V_2.1 was not different between the H-89+forskolin group and the control group (Figure 5G and H and Table S3; control vs. H-89+forskolin: P = 0.3559, Mann-Whitney test). However, the mean NND significantly increased in H-89 treated samples in comparison to two other groups (Figure 5G–H and Table S3; control vs. H-89: P < 0.0001, H-89 vs. H-89+forskolin: P < 0.0001, Mann-Whitney test). Analysis of the mean WPD between Munc13-1 and Ca_V_2.1 gave similar results (Figure 5I and J; Table S3; control vs. H-89+forskolin: P = 0.4810; control vs. H-89: P < 0.0001, H-89 vs. H- 89+forskolin: P < 0.0001, Mann-Whitney test). This is consistent with a decrease in strength of synaptic transmission after H-89 application (Vandael et al., 2020). This change was particularly evident in the cumulative frequency distribution of NNDs and mean WPD between Munc13-1 and Ca_V_2.1 (Figure 5H and 5J; NND – control vs. H- 89: P < 0.0001, control vs. H-89+forskolin: P = 0.3659, H-89 vs. H-89+forskolin: P < 0.0001, Mann-Whitney test; WPD – control vs. H-89+forskolin: P = 0.4810; control vs. H-89: P < 0.0001, H-89 vs. H-89+forskolin: P < 0.0001, Mann-Whitney test). These results indicate that the forskolin-induced changes in Munc13-1 spatial distribution are largely PKA-dependent, implying PKA-mediated changes in vesicle priming.

**Figure 5.**
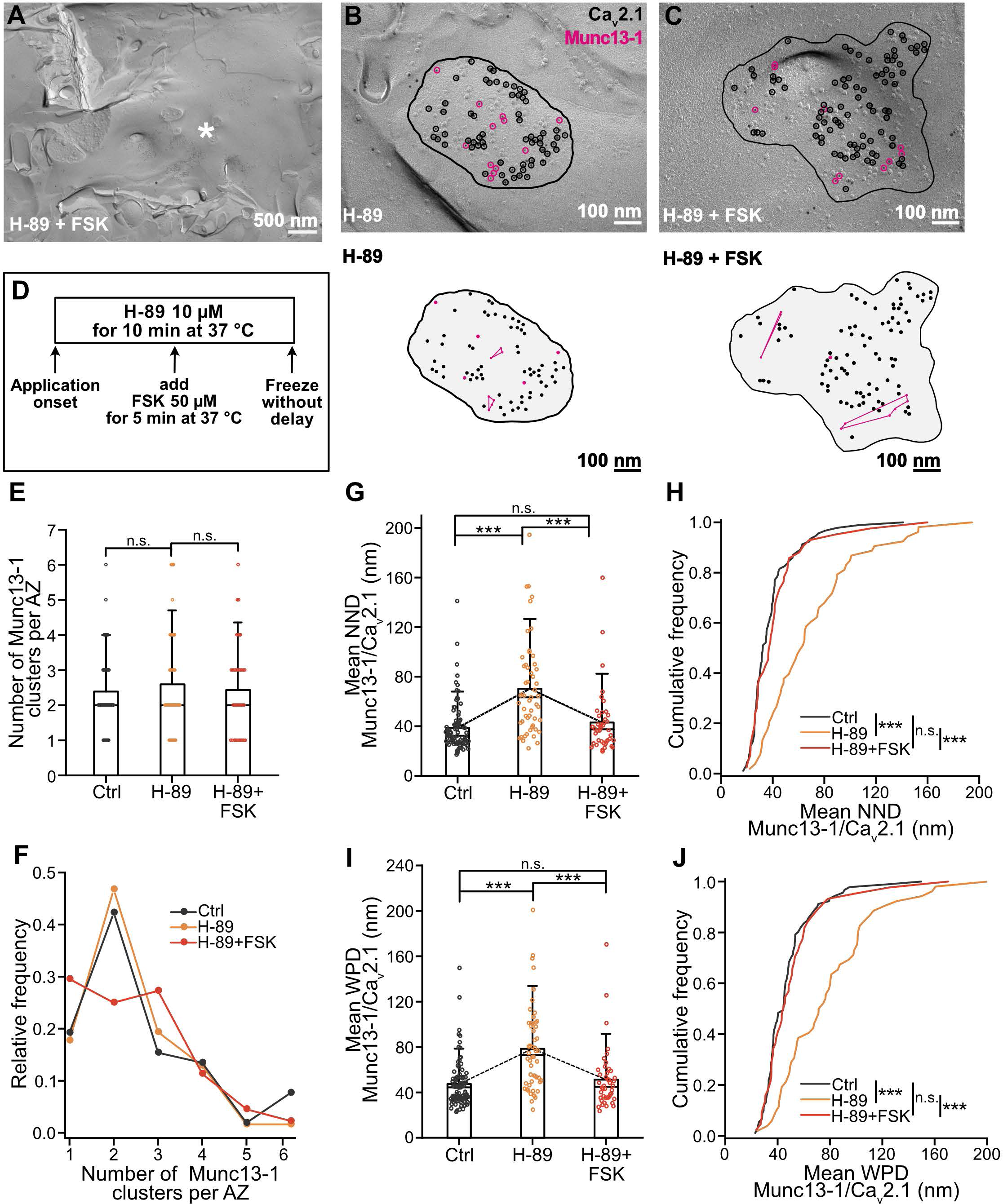
**PKA-dependent remodeling of MFB AZs during chemical potentiation.** **(A)** Example TEM micrographs of freeze-fractured replicas of acute slices showing putative MFB (white asterisk) after H-89 and forskolin treatment (“H-89+FSK”). **(B** and **C)** Top panel: Higher magnification micrographs with putative MFB AZs (black line) co-labeled against Ca_V_2.1 (black empty circles) and Munc13-1 (pink empty circles) in H-89 treated sample (“H-89”, B) and after H-89 with FSK treatment (“H- 89+FSK”, C). Bottom panel: schematic of the above AZs with Munc13-1 clusters (pink circles connected with pink line) in H-89 treated sample (“H-89”, B) and after H-89 with FSK treatment (“H-89+FSK”, C). Note that some Munc13-1 particles do not belong to any cluster and are considered “noise” points (single pink circles). (D) Schematic representation of the time course of the experiment with 10-min H-89 and 5-min forskolin (FSK) treatment. (E) Summary bar graph of the number of Munc13-1 clusters per AZ in DMSO control (“Ctrl”, gray) and after PKA inhibitor H-89 (“H-89”, orange), and H-89 with forskolin (“H- 89 + FSK”, red). Bars and whiskers show mean + SD. Horizontal black lines indicate median values. (F) Relative frequency distribution of data shown in (D), color scheme is identical to (E). (G) Summary bar graph of the mean NND between Munc13-1 and Ca_V_2.1 in DMSO control (“Ctrl”, gray) and after H-89 (“H-89”, orange), and H-89 with FSK treatment (“H- 89 + FSK”, red). Bars and whiskers show mean + SD. Horizontal black lines indicate median values. (H) Cumulative plots of mean NND between experimental Munc13-1 and Ca_V_2.1 point patterns, color scheme is identical to (D). (I) Summary bar graph of the mean WPDs between Munc13-1 and Ca_V_2.1 in DMSO control (“Ctrl”, gray) and after H-89 (“H-89”, orange), and H-89 with FSK treatment (“H- 89 + FSK”, red). Bars and whiskers show mean + SD. Horizontal black lines indicate median values. (J) Cumulative plots of mean WPDS between experimental Munc13-1 and Ca_V_2.1 point patterns, color scheme is identical to (D). Scale bar sizes are indicated on the figure panels.

## DISCUSSION

Using a combined biophysical and ultrastructural analysis, we examined the mechanisms underlying chemical potentiation in a cortical glutamatergic synapse. Functional analysis revealed that chemical potentiation increased both RRP and P_r_, by 146% and 49%, respectively. Structural analysis demonstrated that chemical potentiation increased the number of vesicles and the number of clusters of the priming protein Munc13-1 near the plasma membrane, indicating an increase in the abundance of docked and primed vesicles. Furthermore, we found a significant reorganization of the coupling configuration, including a slight reduction in the NND between presynaptic Ca^2+^ channels and Munc13-1 (i.e. structural coupling). Thus, our results reveal structural correlates of changes in both, RRP and P_r_. Taken together, our results provide new insights into the mechanisms of chemical potentiation at a central synapse.

### Increase in functional and structural vesicle pool size

It is widely assumed that long-term presynaptic plasticity at hippocampal mossy fiber synapses is associated with changes in P_r_ (Castillo, 2012; Monday et al., 2018). In contrast, previous work on the mechanisms of PTP, a short-term form of synaptic plasticity, suggests that primarily changes in pool size are involved (Vandael et al., 2020). Our results indicate that forskolin-induced chemical potentiation also markedly increases vesicle pool size. As forskolin-induced potentiation occludes with LTP at this synapse (Weisskopf et al., 1994), this suggests that both short-term and long-term plasticity converge on similar pool mechanisms. Thus, both short-term and long-term mechanisms may involve the formation of vesicle pool engrams (Vandael et al., 2020).

In addition to the biophysical analysis, our structural data further support a pool engram model. Forskolin increases the size of the RRP by 146%, i.e. to 246% of control value. Structural analysis indicates that the number of docked vesicles per 100 nm profile length changes from 0.78 to 1.20, i.e. to 154%. However, these one-dimensional measurements (per profile length) may need to be converted into two-dimensional predictions (per AZ area). In the extreme case, the number of docked vesicles per AZ might change to (154%)^2^ = 237%. Thus, the increase in the RRP and the increase in number of docked vesicles may be in reasonable agreement. The expansion of the docked vesicle pool may be transient, as Maus et al. (2020) observed only a trend towards an increase in the number of docked vesicles in MFB cryo-fixed with longer delay after the forskolin application. RRP estimation relates to the whole bouton and it is possible that other structural changes can contribute to the increase in RRP, for example increase in AZ area or number (Orlando et al., 2021). In contrast, the increase in Munc13-1 clusters in presynaptic terminals is smaller, from 2.4 to 3.0 per AZ, i.e. to 127%. However, as a caveat, it has to be noted that these values were obtained from both partial and complete AZ replicas. Moreover, it is possible that recently primed Munc13-1 proteins are not as firmly attached to the plasma membrane and might dissociate during the FRL procedure. Alternatively, ubMunc13-2 or Munc13- 3 could be involved in enhanced priming after chemical potentiation (Augustin et al., 1999a).

### Changes in coupling topography during chemical potentiation

Our biophysical analysis reveals that forskolin increases P_r_, although the effects are smaller than those on the vesicular pool. Similar changes in RRP and P_r_ have been reported during PTP (Vandael et al., 2020). Moreover, our results revealed that baseline activity of cAMP-PKA signaling pathway is important for basal synaptic transmission, coupling distance, and sustained activity, as the application of PKA antagonist leads to “uncoupling” of Munc13-1s and Ca_V_2.1 calcium channels and a decrease in synaptic strength (Vandael et al., 2020). However, the mechanisms have remained unclear. One possibility is that an increase in the number of Ca^2+^ channels may lead to increased Ca^2+^ inflow, thereby enhancing transmitter release (Fukaya et al., 2021). Nevertheless, our results show that the number of Ca_V_2.1 channels and Ca_V_2.1 channel clusters remained unchanged. In contrast, forskolin significantly altered the coupling configuration between Ca^2+^ channels and primed vesicles. Under control conditions, we found that the NND between Munc13-1/2 and Ca^2+^ channels under control conditions was 38.8 and 40.3 nm, whereas the mean WPD was 47.3 and 51.6 nm. This is in reasonable agreement with the biophysical coupling distance based on exogenous Ca^2+^ chelator experiments (∼70 nm; Vyleta and Jonas, 2014), considering effects of distant Ca^2+^ channels to transmitter release, contributions of Ca_V_2.2 (N-type) and Ca_V_2.3 (R-type) Ca^2+^ channel subtypes (Castillo et al., 1994; Li et al., 2007), and free rotation of antibodies around the epitope.

In the presence of forskolin, the Munc13-1/ Ca_V_2.1 NND decreased by ∼6 nm. Can this small change explain the increase in P_r_? Although the effect on the NND seems quantitatively small, the effects on transmitter release will be highly nonlinear. Modeling of Ca^2+^ domains showed that tightening of coupling increases P_r_ ∼10-fold per 20 nm change in coupling distance (Bucurenciu et al., 2008). Thus, a 6 nm decrease in the coupling distance may increase P_r_ up to 3 fold. Altogether, our results suggest that chemical potentiation at hippocampal mossy fiber synapses is associated with complex changes in both vesicle pool organization and coupling configuration.

### RRP versus docked and primed pool

The quantitative relation between functional and structural vesicle pools at central synapses has remained controversial. For example, the RRP is often equated with the pool of docked vesicles (Imig et al., 2014), but also may represent a subpool of primed vesicles (Neher and Brose, 2018). Our results shed new light on this aspect. Under control conditions, we estimate an RRP size of 46 vesicles per MFB and 1.62 vesicles per AZ (Table S4; assuming that a typically sized MFB contains 29 AZs; Rollenhagen et al. 2007). Our estimates of the number of docked vesicles per AZ at hippocampal mossy fiber synapses are ∼13 per AZ. In contrast, the putative number of primed vesicles, as determined by the number of Munc13-1 and bMunc13-2 clusters per AZ, is ∼4. Thus, the RRP is in better agreement with the number of Munc13-1 / bMunc13- 2 clusters, presumably corresponding to primed vesicles. Thus, although the RRP, Munc13-1 / bMunc13-2 clusters, and docked vesicle pool correlate, they may represent different functional and structural states of SVs. Consistent with this hypothesis, capacitance measurements from hippocampal mossy fiber terminals suggest that long depolarizing pulses release ∼1400 vesicles, corresponding to 48 vesicles per AZ (Hallermann et al., 2003; Delvendahl et al., 2016). This number is much larger than the number of primed vesicles and even exceeds the number of docked vesicles. Thus, the long depolarizing pulse may trigger the release of loosely docked or tethered vesicles or, in addition, activate refilling.

### Signaling pathways and molecular targets in chemical potentiation

Forskolin increases cAMP levels by direct activation of the AC. Moreover, isoproterenol has effects comparable to forskolin, suggesting that beta 1/2 adrenoreceptors and G_S_ proteins activate the same pathway more physiologically. But what are the molecular events downstream of AC? The effects of forskolin on both the RRP and the primed vesicles are blocked by H-89, suggesting an involvement of PKA. PKA is thought to phosphorylate several presynaptic target proteins, including synaptotagmin 12, tomosyn, rabphilins, SNAP 25 (synaptosomal-associated protein of 25 kDa), and RIMs (Rab3-interacting molecules) (reviewed by Shahoha et al., 2022). For example, phosphorylation of SNAP 25 protein by PKA affected the size of the releasable pool (Nagy et al., 2004). Furthermore, the C_2_A domain of Munc13-1 interacts with the zinc finger domain of RIM building a heterodimeric complex which is critically involved in docking and priming (Lu et al. 2006; Camacho et al., 2017; Zarebidaki et al., 2020). Although PKA dependent phosphorylation of RIM has been shown not be involved in long-term potentiation (Kaeser et al., 2011), recent evidence highlights the importance of other phosphorylation sites of RIM during plasticity (Müller et al., 2022). Thus, the phosphorylation of RIMs may change binding of Munc13 and thereby priming of SVs. In this context, it is interesting that forskolin changes only the localization of Munc13-1, but not bMunc13-2, which lacks C_2_A domain and is thought to prime vesicles independently of RIM (Augustin et al., 1999a; Kawabe et al., 2017). Bassoon, a scaffold protein, is another candidate for phosphorylation, as it strongly colocalizes with Munc13-1 but not Munc13-2 at AZs (Chen et al., 2013). This may suggest that Munc13-1 may be selectively translocated to the protein network of the AZ. However, contributions of other target proteins for PKA-mediated phosphorylation cannot be excluded.

### Neuromodulation of mossy fiber transmission and plasticity

Our results indicate that not only the AC activator forskolin potentiates mossy fiber synaptic transmission. Also, the more natural agonist isoproterenol shows this effect, although its action appears to be more variable (Figure 2). Several neuromodulators, such as acetylcholine (Vogt and Regehr, 2001), dopamine (Kobayashi and Suzuki, 2007), and noradrenaline (Kobayashi et al., 2022) have been reported to affect mossy fiber synaptic transmission. Within the hippocampus, the *stratum lucidum (*the region of mossy fiber termination) is the region with the highest density of innervation by neuromodulatory inputs, such as noradrenaline, adrenaline, and dopamine (Loy et al., 1980; Moudy et al., 1993). Moreover, neuromodulators can change the induction rules and time course of mossy fiber plasticity (Hopkins and Johnston, 1988; Huang and Kandel, 1996). However, whether neuromodulators directly affect the efficacy of synaptic transmission has remained controversial. Our results demonstrate that isoproterenol, a beta 1/2 adrenergic receptor agonist, can directly potentiate synaptic transmission in the subset of MFB at the unitary level.

### Implications for hippocampal network function

Our findings indicate that neuromodulation can regulate single-synapse computations at the hippocampal mossy fiber synapse (Abbott and Regehr, 2004; Silver, 2010). The unitary EPSP recordings (Figure 2) provide direct evidence for this hypothesis, demonstrating that chemical potentiation can switch the hippocampal mossy fiber synapse from a subdetonation into a detonation mode (Henze et al., 2002; Vyleta et al., 2016). Previous work showed that facilitation and PTP may lead to conditional detonation or plasticity-dependent detonation at this synapse. Our results suggest that neuromodulation can have similar effects. Thus, burst or superburst activity in GCs (Zhang et al., 2020) may lead to detonation, but sparse activation of GCs in conjunction with activation of neuromodulatory pathways may also efficiently activate the CA3 network. Notably, stimulation of locus coeruleus neurons, presumably releasing noradrenaline or dopamine, enhances hippocampus-dependent memory (Takeuchi et al., 2016; Kempadoo et al., 2016; Wagatsuma et al. 2018; Kaufman et al. 2020). It has been suggested that noradrenaline release from locus coeruleus neurons may signal arousal, novelty, and reward prediction (Sara et al., 2009). However, the mechanisms by which this enhances memory remain unclear. Our results suggest that neuromodulator-dependent strengthening of the hippocampal teacher synapse could be involved in these complex effects. Overall, the phenomenal similarities between chemically and HFS-induced potentiation suggest a common mechanism behind MFB plasticity. Future work should determine whether these presynaptic mechanisms are conserved across synapses, brain regions, and species.

## LEAD CONTACT AND MATERIALS AVAILABILITY

Further information and requests for resources, original data, and analysis programs reagents should be directed to and will be fulfilled by the Lead Contact, Peter Jonas upon reasonable request. This study did not generate new unique reagents. Computer code was not generated in this study.

## ACKNOWLEDGEMENTS

We thank Carolina Borges-Merjane, Jing-Jing Chen, Katharina Lichter, and Samuel Young for critically reading the manuscript; the Electron Microscopy Facility of ISTA, in particular Vanessa Zheden and Walter Kaufmann for extensive support, advice, and experimental assistance; the Preclinical Facility of ISTA, in particular Victoria Wimmer and Michael Schunn, for experimental assistance; Florian Marr and Christina Altmutter for technical support; Alois Schlögl for help with analysis; and Eleftheria Kralli-Beller for manuscript editing. We also thank Cordelia Imig for generating and kindly providing us Munc13-1cKO-Munc13-2/3^(−/−)^ animals. This project received funding from the European Research Council and European Union’s Horizon 2020 research and innovation programme (ERC 692692) and from Fond zur Förderung der Wissenschaftlichen Forschung (Z 312-B27 Wittgenstein award; W1205-B09, P36232- B), both to P.J. Part of the work has been published in O.K.’s thesis in partial fulfillment of the requirements for the degree of Doctor of Philosophy.

## AUTHOR CONTRIBUTIONS

O.K. (EM) and Y.O. (electrophysiology) performed all experiments and data analysis; O.K., Y.O., R.S. and P.J. conceived the study and designed experiments; N.B. provided resources; O.K. and P.J. wrote the paper. All authors jointly revised the paper.

## .METHODS

### Animal experiments

Animals were bred in a colony maintained in the Preclinical Facility at ISTA. All procedures strictly complied with institutional, Austrian, and European ethical regulations for animal experiments, and were approved by the Bundesministerium für Bildung, Wissenschaft und Forschung of Austria (Reference number 2020- 0.648.587 and 2022-0121.440).

To determine antibody specificity against Munc13-1 and bMunc13-2, a Tg(Prox1-Cre)SJ32Gsat/Mmucd line (Mutant Mouse Resource & Research Centers) was crossed with mice in which a floxed Munc13-1 was inserted on a Munc13-2/3^(−/−)^ background (Hilton et al., 2022; Banerjee et al., 2022). Mice were genotyped, DNA extracted from toe or ear clippings, to ensure homozygosity of progenies.

### Slice preparation

Acute slices from 21–40-day-old (P21–40) C57BL/6 mice and 21–25-day-old (P21– 25) Wistar rats were used. Mice were used for all EM and FRL experiments as an established preparation for HPF (Borges-Merjane et al., 2020). Wistar rats were used for paired electrophysiological recordings based on previously established protocols for optimal preservation of hippocampal mossy fiber tract (Bischofberger et al., 2006b). Mice or rats were lightly anesthetized with isoflurane and rapidly decapitated. The brain was dissected from the skull and blocking “magic-cut” was performed under ice-cold high-sucrose solution containing: 87 mM NaCl for mice and 64 mM NaCl for rats experiments, 75 mM sucrose for mice or 120 mM for rats, 25 mM NaHCO_3_, 10 mM D-glucose, 2.5 mM KCl, 1.25 mM NaH_2_PO_4_, 0.5 mM CaCl_2_, and 7 mM MgCl_2_, equilibrated with 5% CO_2_/95% O_2_, osmolarity ∼325 mOsm. Transverse hippocampal slices were sectioned at a thickness of 150 μm for HPF or 350 μm for paired electrophysiological recordings, using a vibratome (VT1200S, Leica Microsystems) in ice-cold high-sucrose solution. Slices were transferred to a maintenance chamber and recovered at 35°C for 30–45 min. After recovery and until use for electrophysiology recordings, slices were kept in the maintenance chamber with a high-sucrose solution at room temperature. For HPF experiments, slices were kept until freezing in a second set of maintenance chambers filled with artificial cerebrospinal fluid (ACSF) solution, identical to solution used for electrophysiology recordings, containing: 125 mM NaCl, 25 mM D-glucose, 25 mM NaHCO_3_, 2.5 mM KCl, 1.25 mM NaH_2_PO_4_, 2 mM CaCl_2_, and 1 mM MgCl_2_, equilibrated with 5% CO_2_/95% O_2_.

### Paired recordings

Slices were placed in a recording chamber and superfused with ACSF for at least 15– 30 minutes before onset of recording. Subcellular patch-clamp recordings from single MFBs and simultaneous recordings from CA3 PNs were performed under an upright microscope with infrared difference interference contrast as previously described (Vandael et al., 2021). Pre- and postsynaptic recording pipettes were fabricated from borosilicate glass capillaries (2.0 mm outer diameter, 1.0 mm inner diameter), and had open-tip resistances of 10–25 MΩ and 2–5 MΩ, respectively, when filled with a K^+^-based internal solution (120 mM K-gluconate, 20 mM KCl, 2 mM MgCl_2_, 2 mM Na_2_ATP, 10 mM HEPES, and 10 mM EGTA, pH adjusted to 7.3 with KOH, ∼300 mOsm). The pre- and postsynaptic holding potential was set at −70 mV under voltage-clamp configuration. APs in MFBs were evoked by brief voltage pulses (amplitude 800–900 mV, duration 0.1 ms) in the tight-seal, cell-attached configuration. A train stimulation (10 stimuli at 50 Hz) was delivered once every 20 s (i.e. at 0.05 Hz). Postsynaptic series resistance was 9.03 ± 0.08 MΩ (mean ± SEM; median 8.62 MΩ; ranging from 5.08 to 14.84 MΩ; 18 pairs). Series resistance was uncompensated, but carefully monitored with a test pulse (5 mV) following each data acquisition sweep. Current-clamp recordings were performed at approximately −70 mV with <150 pA injection of hyperpolarizing current. A 50-Hz train stimulation (3 pulses) was delivered once every 20 s. Membrane potential was checked repeatedly throughout the experiment, and holding current was carefully readjusted if required. In current-clamp mode, pipette capacitance was ∼70% compensated and series resistance was balanced using the bridge circuit of the amplifier. All electrophysiology recordings were performed at room temperature (22–25°C).

### Chemical potentiation experiments

To induce chemical potentiation in MFB synapses, 50 µM forskolin (Tocris Bioscience) in 0.01% dimethyl sulfoxide (DMSO; Sigma Aldrich) was applied to ACSF solution, slices were incubated for 5 min at ∼37°C (HPF) or room temperature (22–25°C; electrophysiology). Acute slices were frozen immediately after the onset of forskolin application (HPF). To activate of G_s_-coupled receptors, 1 µM of isoproterenol hydrochloride (Tocris Bioscience) or (−)-Isoproterenol hydrochlorid (Sigma-Aldrich) were used, slices were incubated for 10 min at room temperature (electrophysiology). During electrophysiology recordings, forskolin and isoproterenol were applied by bath application at the specified concentration. Perfusion rate was 4–5 ml min^−1^. During the recording of mEPSCs, 1 µM tetrodotoxin (TTX; Alomone Labs) and 10 µM gabazine (BioTrend) were added directly to ACSF.

To block PKA activity in MFB synapses, samples were incubated in H-89- containing ACSF (10 µM in 0.01% DMSO, Tocris Bioscience) for 10 min prior to freezing.

### HPF on acute hippocampal slices

HPF was performed with a Leica EM ICE high-pressure freezing apparatus (Leica Microsystems) as previously described (Borges-Merjane et al., 2020). Materials and samples were always kept close to physiological temperature (∼37°C). After slicing and recovery procedures, hippocampal acute slices were frozen with filler medium containing 15% polyvinylpyrrolidone (PVP; Sigma Aldrich) in ACSF, equilibrated with 5% CO_2_/95% O_2_ gas mixture, and kept at 37°C. The specimen sandwich for HPF was assembled with two 120-μm sapphire discs, a 150-µm thick spacer ring, a 450-μm thick top ring (Engineering Office M. Wohlwend, Sennwald, Switzerland), and transparent half-cylinder cartridges and middle-plate (Leica Microsystems). The outer diameter of sapphire disks and spacer rings was 6 mm. The inner diameter of the rings was either 5 mm or 4 mm depending on the size of the slice.

### Freeze-substitution and ultramicrotomy

For the freeze-substitution experiments (Borges-Merjane et al., 2020), the HPF samples were put to vials with 0.1% tannic acid (Sigma Aldrich) in acetone, transferred to a Leica AFS2, kept at −90°C, and shaken for 20–22 hours. Next, samples were washed 3–4 times 10 min each with fresh acetone chilled to −90°C and left at the same temperature for 6 hours in acetone containing 2% osmium (Science services (EMS)) and 0.2% uranyl acetate (Serva). Subsequently, the temperature was raised to −60°C (10°C per hour) and was kept at −60°C for 3 hours. Then samples were heated to −30°C (10°C per hour), and kept at −30°C for 3 hours; then the temperature was finally raised to 0°C (10°C per hour). Next, the vials were washed with ice-cold acetone, 3 times for 10 min each, on ice. Then they were washed twice with propylene oxide for 10 min each on ice, and infiltrated with hard Durcupan resin (11.4 g reagent A, 10 g B, 0.3 g C, and 0.1 g D; all Sigma-Aldrich) at 2:1, 1:1, and 1:2 propylene oxide/Durcupan resin mix for at least 1 hour each, shaking at room temperature. Samples were left in pure resin overnight. Samples were polymerized overnight at 100°C in BEEM capsules. 70 nm ultrathin sections were cut with a Leica EM UC7 ultramicrotome with diamond knives (Diatome Histo). Sections were picked up on Formvar-coated copper slot grids for transmission electron microscopy (TEM) imaging. Post-staining was done with 4% uranyl acetate in water for 10 minutes, followed by lead citrate for 2–3 minutes (Sigma-Aldrich).

### HPF for freeze-fracture experiments

For evaluation of replicas from acute hippocampal slices, the tissue was prepared as described above with some modifications. For these experiments, 4.6-mm gold-plated carriers with 140-µm double-sided adhesive tape were used. Freeze fracture replicas were produced using the tensile fracture approach with two freeze-fracture machines interchangeably: JFD V (Jeol) and ACE 900 (Leica Microsystems). Samples were fractured at −120°C under a high vacuum. Carbon-platinum replicas were prepared by evaporating layered carbon (C) at a 90° angle and platinum (Pt) at 60° on the surfaces of the slices: the first C layer was 5 nm thick, the second Pt layer was 2 nm, and the final third C layer was 20 nm. Afterwards, replicas were brought to Tris-buffer solution (TBS, 50 mM, pH 7.4) at room temperature and transferred to glass tubes containing 2.5% sodium dodecyl sulfate (SDS) solution and 20% sucrose in 15 mM TBS (pH 8.3). The replicas were solubilized in SDS solution for 18 hours, shaking at 80°C. Next, prepared replicas were kept in the same SDS solution until immunolabeling experiments.

### Freeze-fracture replica immunolabeling (FRL)

All replicas were handled with utmost care during all steps to ensure they stayed intact. Washes were done in 12-well porcelain plates and replicas were transferred with a glass rod. First, replicas were washed with fresh 2.5% SDS in TBS solution for 10 min shaking at room temperature. Then they were transferred to 0.1% Tween in 0.05% bovine serum albumin (BSA, Sigma Aldrich) in TBS buffer solution and washed for 10 min. Afterwards, sections were washed 3-4 times for 15 min in 0.05% BSA in TBS, before blocking for 90 min in 5% BSA. Then, replicas were incubated in the first primary antibody, guinea pig anti-Ca_V_2.1 (P/Q-type; Synaptic Systems, Cat # 152 205, RRID:AB_2619842, 1.3 µg ml^−1^; Chen et al., 2023) in 2% BSA in TBS overnight, shaking at 15°C. Next, replicas were washed 3–4 times in 0.05% BSA in TBS for 15 min each and again blocked with 5% BSA in TBS for 90 min. Replicas were incubated in second primary antibodies, rabbit anti-Munc13-1 (Synaptic Systems, Cat # 126 103, RRID:AB_887733) or anti-Munc13-2 (Synaptic Systems, Cat # 126 203, RRID:AB_2619807; each 2.5 µg ml^−1^) overnight, shaking at 15°C. The Munc13-2 antibody was targeted against the brain-specific splice variant bMunc13-2, which is more abundant than the ubiquitous splice variant ubMunc13-2 in the hippocampus (Breustedt et al., 2010; Kawabe et al., 2017). Then, the same washing and blocking steps were repeated. Secondary antibodies goat anti-rabbit 5-nm gold conjugated (BBI Solutions, Cat # EM GAR5, RRID:AB_1769142) and goat anti-guinea pig 10-nm gold conjugated (BBI Solutions, Cat # EM.GAG10, RRID:AB_2892072) at concentration 1:30 in 2% BSA in TBS were sequentially applied overnight at 15°C. Sections were picked up on Formvar-coated copper mesh grids for TEM imaging. For unequivocal distinction of secondary antibodies in case of double labeling, two sizes of gold-conjugated particles were used – 5 and 10 nm. Batches of antibodies were tested and distribution of the particle sizes showed a clear separation between two populations (mean size of 5-nm antibody: (mean ± SEM) 5.07 ± 0.07 nm; 10-nm antibody: 10.12 ± 0.08 nm). The specificity of anti-Munc13-1 and anti-bMunc13-2 antibodies was confirmed using replicas of acute hippocampal slices prepared from Munc13-1cKO- Munc13-2/3^(−/−)^ animals (Hilton et al., 2022; Banerjee et al., 2022). The number of particles per 0.1 µm^2^ AZ of MFB from knock-out (KO) animals was compared to wild-type (WT) data from acute hippocampal slices cryo-fixed without DMSO (anti-Munc13- 1 WT: 18.9 ± 1.5 (mean ± SEM), median 17.5, n = 40 AZ, N = 3 mice; cKO: 0.5 ± 0.1, median 0, n = 77 AZ, N = 3 mice; WT vs. cKO: P < 0.0001, Mann-Whitney test; anti- bMunc13-2: WT: 15.2 ± 1.5, median 13.6, n = 40 AZ, N = 3 mice; KO: 0.5 ± 0.2, median 0, n = 53 AZ, N = 3 mice; WT vs. KO: P < 0.0001, Mann-Whitney test).

## QUANTIFICATION AND STATISTICAL ANALYSIS

### Electrophysiology data analysis

Data were acquired with a Multiclamp 700B amplifier, low-pass filtered at 10 kHz, and digitized at 40 or 50 kHz using a CED 1401 plus or power1401 mkII interface (Cambridge Electronic Design, Cambridge, UK). Pulse generation and data acquisition were performed using FPulse version 3.3.3 or 3.45 (U. Fröbe, Freiburg, Germany) under running Igor Pro version 6.37 (Wavemetrics). Data were analyzed with Stimfit version 0.15 and Igor Pro. Only recordings with < 15 MΩ postsynaptic series resistance and stationary postsynaptic responses (EPSC_1_; based on Pearson correlation coefficient; p > 0.05) were included for analysis. During the 50-Hz train stimulation, the peak amplitude of each EPSC was determined as the difference between the mean of current 2-ms preceding the onset of voltage pulse and the mean of current over a time window ± 0.5 ms around the peak. The time window of peak amplitude detection was set between the offset of voltage pulse and the next onset of voltage pulse (19.9 ms). The paired-pulse ratio (EPSC_2_ / EPSC_1_) was calculated from average EPSC amplitudes before (10 individual traces) and after forskolin or isoproterenol application (5 or 10 individual traces).

To estimate RRP size, P_r_, and refilling rate, cumulative EPSC peak amplitudes were plotted against time for 10 stimuli at 50 Hz, the last 4 data points were fit by linear regression, and back-extrapolated to the time point 0 (Schneggenburger et al., 1999). The size of the RRP was determined as the intersection of the regression line with the ordinate. P_r_ was measured as the ratio of EPSC_1_ amplitude over RRP size, and refilling rate was obtained from the slope of the line. Note that the RRP estimates by this method represent “pool decrement” rather than absolute pool size (Neher, 2015).

mEPSCs were detected using a template-fit analysis (Jonas et al., 1993; Clements and Bekkers, 1997) under running MATLAB R2017a (MathWorks).

Membrane potentials are given without correction for liquid junction potentials. In total, data reported in this paper were obtained from 25 MFB–CA3 PN paired recordings and 11 CA3 PNs.

### TEM imaging, AZ profile and FRL analysis

All EM micrographs from ultrathin sections were analyzed blindly towards the condition of treatment. Hippocampal MFBs were identified in the hippocampal CA3b/c subregions in *stratum lucidum*. They were recognized based on previously well-characterized morphological features: large size, high density of clear SVs, presence of large dense-core vesicles, high density of mitochondria, multiple synaptic contacts with large spines, and nonsynaptic *puncta adhaerentia* contacts with dendritic shafts (Studer et al., 2014; Borges-Merjane at al., 2020). AZs were defined as the part of presynaptic membrane directly opposed to the electron dense region of the postsynaptic membrane (postsynaptic density), with an accumulation of clear and round vesicles in close proximity to the membrane and characteristic widening of the synaptic cleft (Rollenhagen et al., 2007; Borges-Merjane at al., 2020). Ultrastructure analysis focused on identifying the number and diameter of vesicles docked at identified AZ profiles.

Micrographs of ultrathin sections were taken with a transmission electron microscope (Thermo Fisher/FEI Tecnai 10, 80 kV acceleration voltage) with an OSIS Megaview III G3 camera and Radius acquisition software. All EM images were analyzed with Fiji open source software (Schindelin at al., 2012). To optimize double membrane visualizations and accurate measurements, brightness and contrast were adjusted in Fiji. Multiple AZ profiles were distinctly analyzed per mouse, at least 3 mice. Vesicles, whose outer membrane and presynaptic AZ membrane were in direct contact, were considered “docked”. For quantitative comparison between groups, numbers of docked vesicles per profile were normalized to 100 nm of AZ profile length. Replicas were imaged as described for ultrathin sections. Similar to the ultrathin sections, MFBs were observed in *stratum lucidum* of the CA3b/c regions, along the mossy fiber tract. In replicas MFBs were identified based on their location, size of the terminals, availability of attached cross fracture with numerous SVs, and several putative AZs on the P-face of plasma membrane (Hagiwara et al., 2005). ImageJ Fiji open-source software was used for the analysis of all replica micrographs. AZs were recognized based on the P-face location, distribution of intramembrane particles (IMPs), and labeling of AZ proteins and Ca^2+^ channels. Gold particles and AZ area were manually segmented. Their corresponding coordinates (i.e. pixel positions on a image) were used for the point-pattern analysis. The number of gold particles per AZ was specified per 0.1 µm^2^ of AZ area and used as a labeling density criteria for direct comparison across groups. NND for single point pattern was calculated as a Euclidian distance from each point to its nearest neighbor. NND between two distinct point patterns within same AZ was calculated as Euclidean distance from each point in the first point pattern (Munc13-1 or 2) to the nearest point in the second point pattern (Ca^2+^ channels). Random NND distributions were calculated from randomly distributed Poisson point patterns with densities identical to experimental values. Mean NNDs were calculated as mean value of all obtained NNDs within or between point patterns. The mean WPD was calculated from each point in the first point pattern (Munc13-1 or 2) to each point in the second point pattern (Ca^2+^ channels). Then, the distances were weighted using following function:

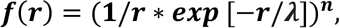

where *r* is measured experimental distance, n is power coefficient (1.77; Vyleta and Jonas, 2014), *λ* is the length constant that depends on fast Ca^2+^ buffer concentration, its kinetics, and resting Ca^2+^ concentration:

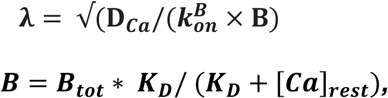

where *D_Ca_* – diffusion coefficient of free Ca^2+^, 220 µm^-2^ s^-1^; *k*^*B*^_*on*_ – rate constant of Ca^2+^ binding to fast exogenous buffer, 4 × 10^8^ M^−1^ s^−1^; *B_tot_* – total Ca^2+^ buffer concentration, 300 µM; *B*, free Ca^2+^ buffer concentration; *K_D_* – dissociation constant of exogenous buffer, 0.22 µM; *[Ca]_rest_ –* resting Ca^2+^ concentration, 0.074 µM (Jackson and Redman, 2003). To compute the weighted distance, pairwise distances from same AZs were multiplied with the corresponding weight factor and averaged. For comparison across groups, weighted distance values were pooled.

Density-based spatial clustering of applications with noise (DBSCAN; Ester et al., 1996) was performed to determine the number of clusters of Ca^2+^ channels (Ca_V_2.1) and Munc13s. The minimum number of points per cluster was set to 2. The size of neighborhood *ε*, the distance between two points that can be part of the same cluster, was determined from “knee” value of 2-nearest neighbor plot (K-plot) for each analyzed AZ. The most *ε* values were in the range of mean ± 2 SD of NND. The mean *ε* values were 37.2 nm (Ca_V_2.1 control), 37.6 nm (Ca_V_2.1 forskolin), 41.3 nm (Munc13- 1 control), 42.2 nm (Munc13-1 forskolin), 54.3 nm (bMunc13-2 control), and 51.2 nm (bMunc13-2 forskolin). The intercluster distance was determined as minimal distances between cluster edges. The analysis of NND and clustering was done in R (RStudio) using spatstat (2.0-1) and dbscan (1.1-8) CRAN packages. The weighting of pairwise distances was done in MATLAB R2017a (MathWorks).

Univariate Ripley K-function (Ripley, 1977) and corrected H-function (Kiskowski et al., 2009) for Ca_V_2.1 and Munc13-1/2 particles were computed using a Matlab software package (Rebola et al., 2019). Mean experimental H-function was compared to mean simulated H-function using maximum absolute difference (MAD) test. Mean simulated H-function was estimated from population of all Monte-Carlo simulated H-functions (100 for each AZ).

### Statistical analysis

Statistical analysis was performed with Origin 2019 (Origin Lab), Prism (GraphPad), and RStudio. All data groups were tested for normality with the D’Agostino-Pearson test, followed by a non-parametric Kruskal-Wallis test. Data groups were then compared using two-sided paired non-parametric Wilcoxon signed-rank or two-sided unpaired non-parametric Mann-Whitney test. For the graphical representation of statistics, * indicates P < 0.05, ** P < 0.01, *** P < 0.001, and **** P < 0.0001. In text and figures, values report mean and median, and errors report standard error of the mean (SEM; paired recordings) or standard deviation (SD; EM and FRL), as specifically stated.

---

**Figure S1.**
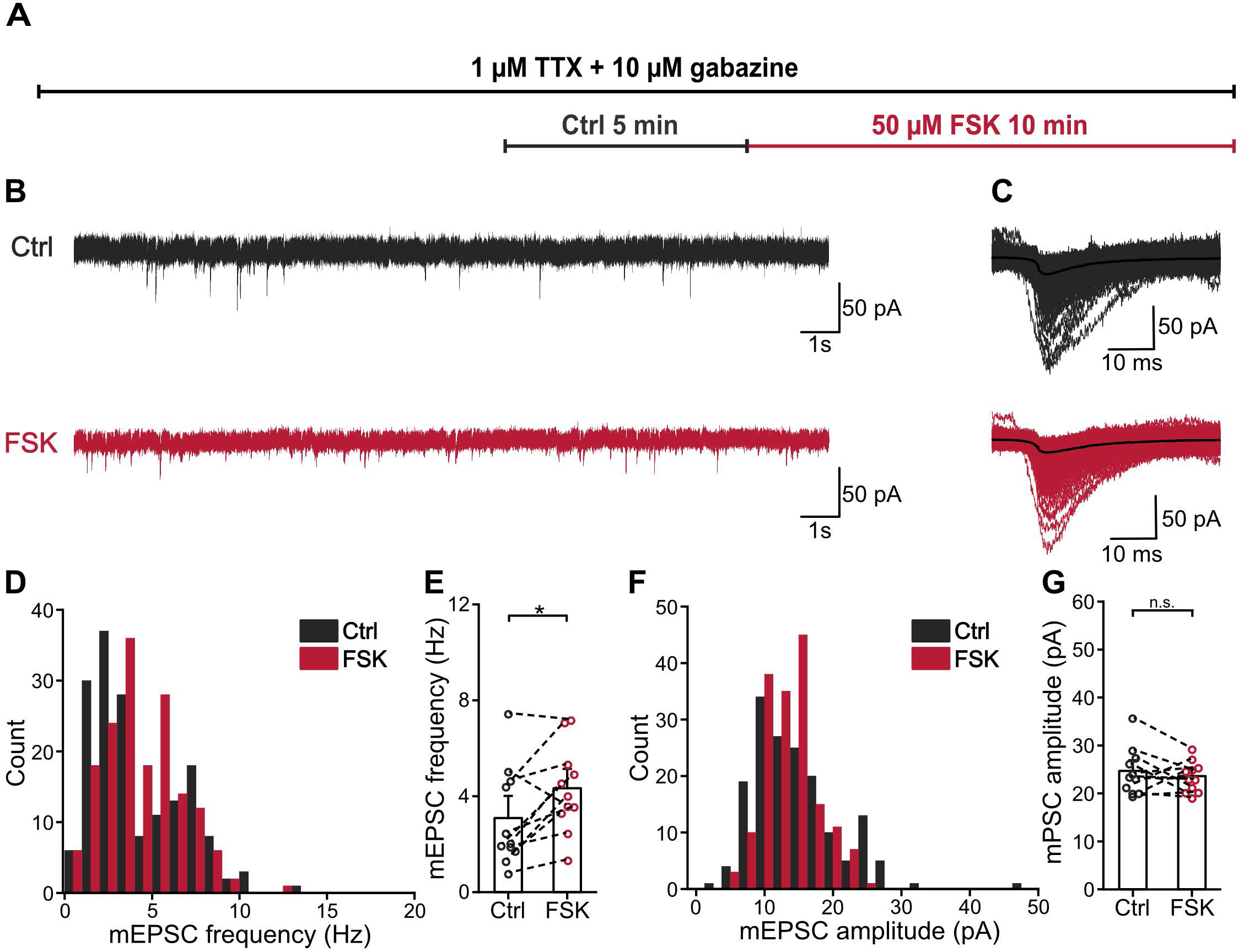
**related to Figure u1. The effect of forskolin on mEPSCs in CA3 PNs.** **(A)** Schematic representation of time course of the experiment. 1 µM TTX and 10 µM gabazine were added to ACSF and perfused for at least 10 min prior to the onset of recordings. Control data (“Ctrl”, gray) was recorded for 5 min prior to forskolin application. Forskolin data was recorded during last 5 min of 10-min forskolin treatment (“FSK”, red). **(B)** Representative traces of miniature EPSCs (mEPSCs) before (“Ctrl”, gray) and after 50 µM forskolin (“FSK”, red). **(C)** mEPSCs at expanded time scale after detection and alignment to the onset time point before (“Ctrl”, gray) and after 50 µM forskolin (“FSK”, red). Black line represents average. **(D)** Histogram of mEPSC frequency before (“Ctrl”, gray) and after 50 µM forskolin (“FSK”, red). Data from 11 cells and 3 rats. **(E)** Summary bar graph of mEPSC frequencies (C) before (“Ctrl”, gray) and after 50 μM forskolin (“FSK”, red). Bars and whiskers show mean + SEM. **(F)** Histogram of mEPSC peak amplitude before and after 50 µM forskolin, color scheme is identical to (D). Data from 11 cells and 3 rats. **(G)** Summary bar graph of mEPSC peak before (“Ctrl”, gray) and after 50 μM forskolin (“FSK”, red). Bars and whiskers show mean + SEM.

**Figure S2.**
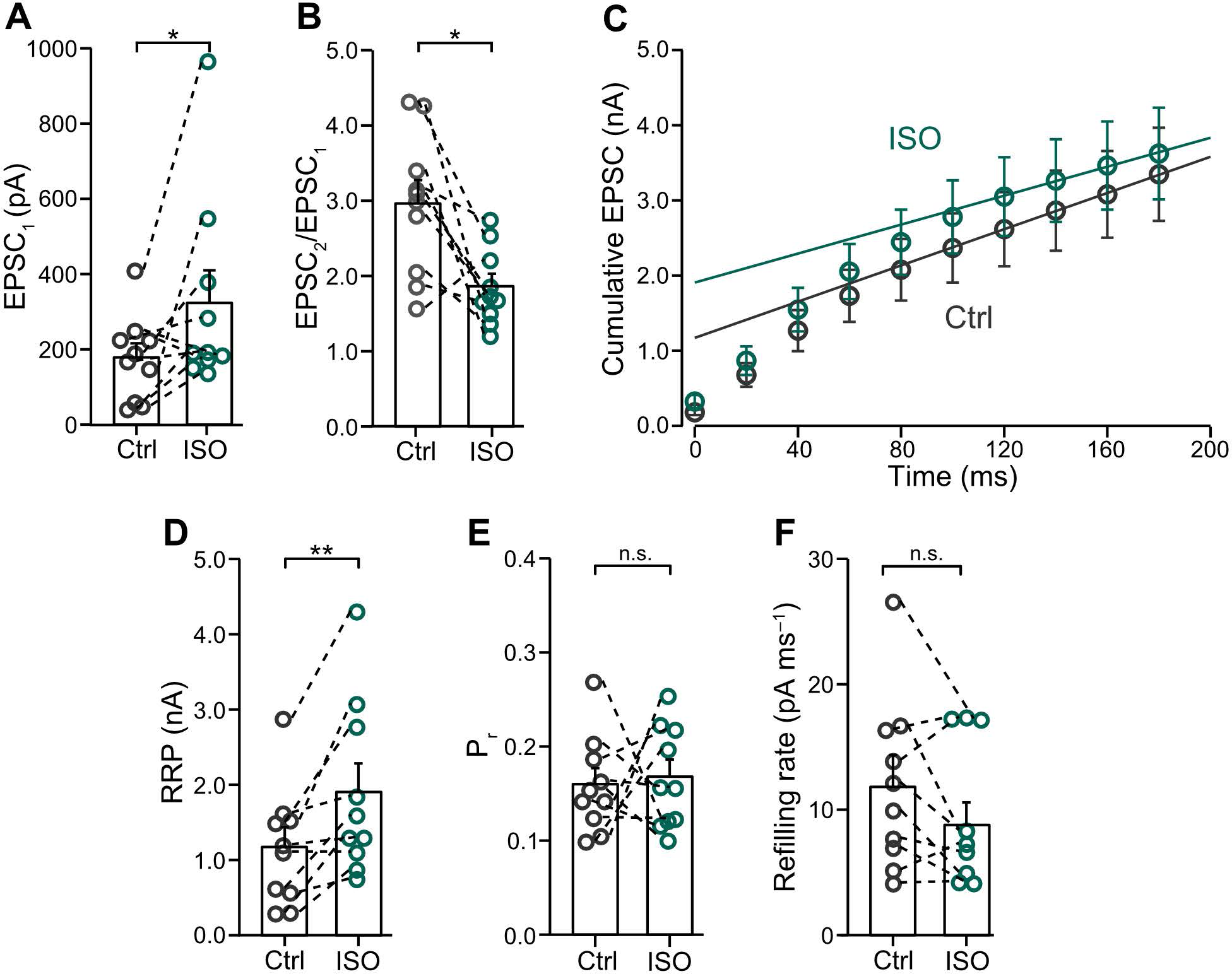
related to Figure 2**. The effect of isoproterenol on MFBs to CA3 PNs synapses.** (A) Summary bar graph of EPSC_1_ peak amplitudes before (“Ctrl”, gray) and in the presence of 1 µM isoproterenol (“ISO”, green). Bars and whiskers show mean + SEM. (B) Summary bar graph of paired-pulse ratio (EPSC_2_ / EPSC_1_) before (“Ctrl”, gray) and in the presence of 1 µM isoproterenol (“ISO”, green). Bars and whiskers show mean + SEM. (C) Cumulative plot of EPSC peak amplitudes during a 50-Hz train with 10 stimuli before (“Ctrl”, gray) and in the presence of 1 µM isoproterenol (“ISO”, green). Data points during the last 4 stimuli (at time points ≥ 120 ms) were fit by linear regression and back-extrapolated to time point 0. **(D–F)** Summary bar graphs of RRP size (D), P_r_ (E), and refilling rate (F), estimated from the cumulative EPSC plot (C), before (“Ctrl”, gray) and in the presence of 1 µM isoproterenol (“ISO”, green). Bars and whiskers show mean + SEM.

**Figure S3.**
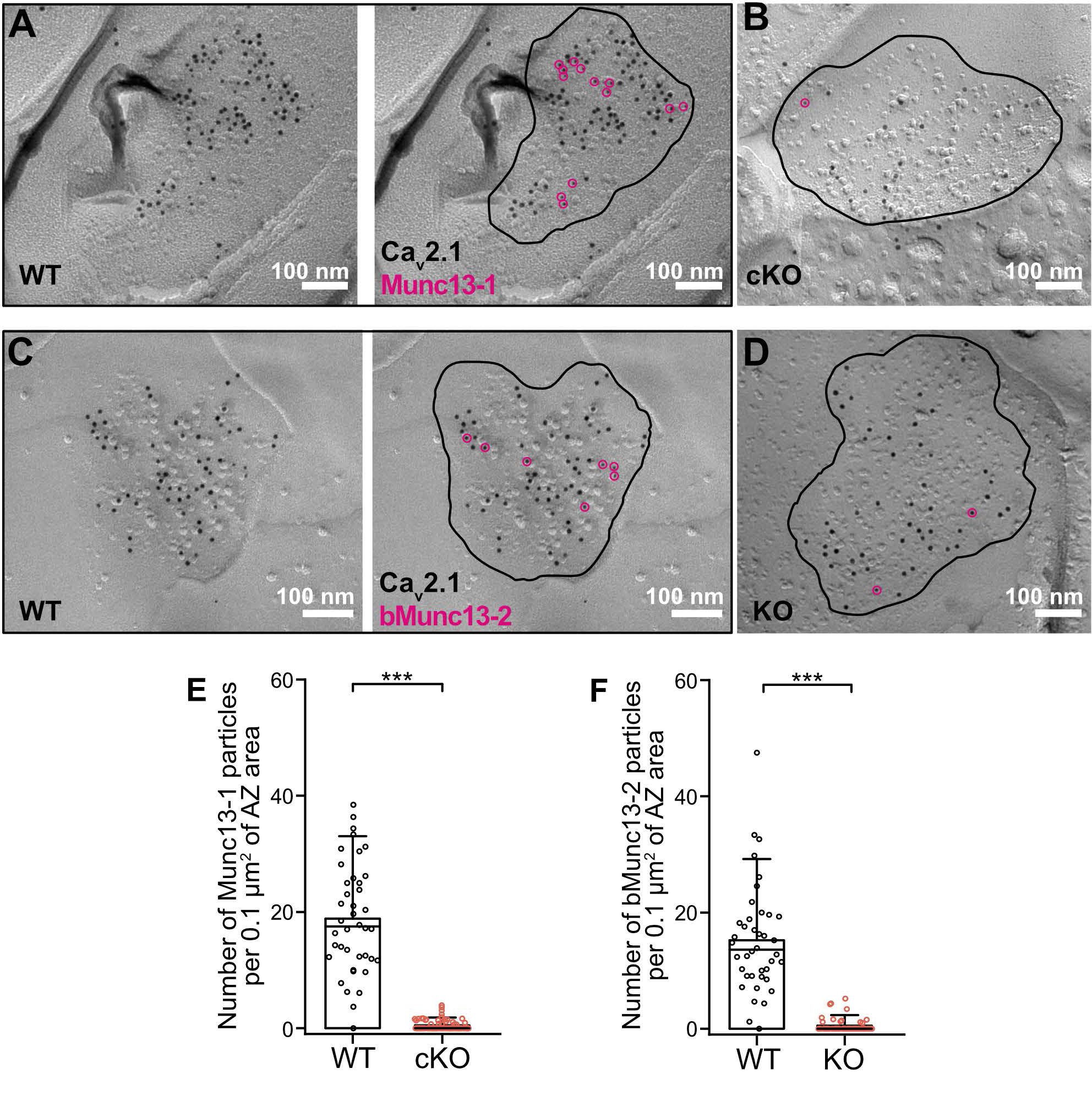
related to Figure 4**. Labeling of Munc13s in MFBs in wild-type and knock-out mice.** (A) Example TEM micrographs of freeze-fractured replicas of acute wild-type (“WT”) slices. Left panel: putative MFB AZ with gold particles of two sizes, 10 and 5 nm respectively. Right panel: putative MFB AZ (black line) co-labeled against Ca_V_2.1 (black) and Munc13-1 (pink empty circles). (B) Example TEM micrographs of freeze-fractured replicas of acute slices from floxed Munc13-1-Prox1Cre mice (“cKO”) showing putative MFB AZ (black line) co-labeled against Ca_V_2.1 (black) and Munc13-1 (pink empty circle). (C) Example TEM micrographs of freeze-fractured replicas of acute wild-type (“WT”) slices. Left panel: putative MFB AZ with 10- and 5–nm gold particles. Right panel: putative MFB AZ (black line) co-labeled against Ca_V_2.1 (black) and bMunc13-2 (pink empty circles). (D) Example TEM micrographs of freeze-fractured replicas of acute slices from Munc13-2/3^(-/-)^ mice (“KO”) showing putative MFB AZ (black line) co-labeled against Ca_V_2.1 (black) and bMunc13-2 (pink empty circles). **(E–F)** Scatter plot of the number of Munc13-1 (E) and bMunc13-2 (F) particles per 0.1 µm^2^ of AZ area in WT control (“WT”, black) and in floxed Munc13-1-Prox1Cre mice (“cKO”, orange) and Munc13-2/3^(-/-)^ mice (“KO”, orange). Bars and whiskers show mean + SD. Horizontal black lines indicate median values. Scale bar sizes are indicated on the figure panels.

**Figure S4.**
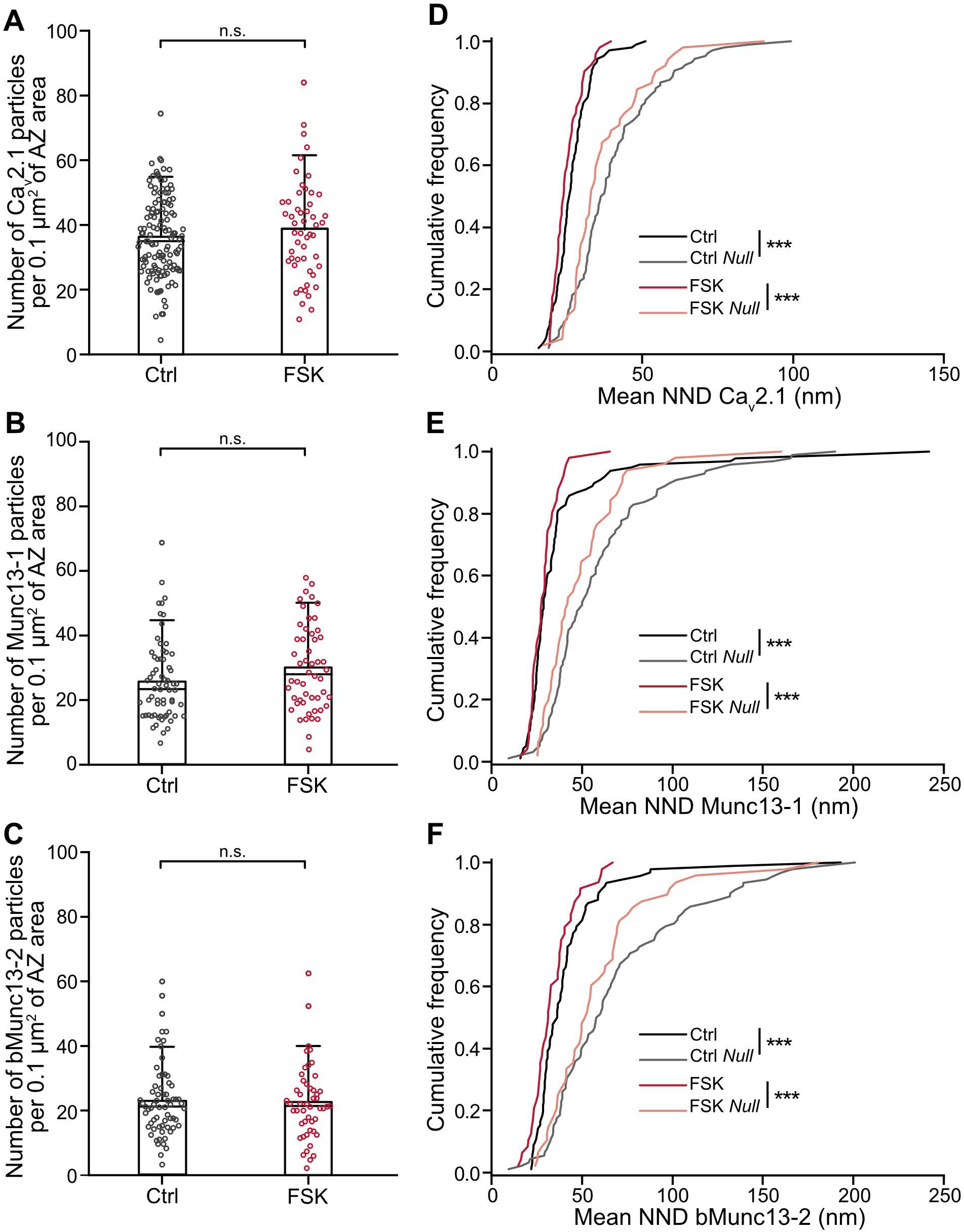
related to Figure 4**. Labeling of Ca_v_2.1s and Munc13s in MFBs before and after 5-min forskolin treatment.** **(A–C)** Scatter plot of the number of Ca_v_2.1 (A), Munc13-1 (B) and bMunc13-2 (C) particles per 0.1 µm^2^ of AZ area in DMSO control (“Ctrl”, gray) and after forskolin (“FSK”, red). Bars and whiskers show mean + SD. Horizontal black lines indicate median values. **(D–F)** Cumulative plots of mean NND between experimental Ca_v_2.1 (D), Munc13-1 (E) and bMunc13-2 (F) point patterns and randomly simulated data in treated groups. Experimental data DMSO control (“Ctrl”, dark gray) and after forskolin (“FSK”, dark red), randomly simulated data DMSO control (“Ctrl *Null”*, light gray) and after forskolin (“FSK *Null”*, light pink).

**Figure S5.**
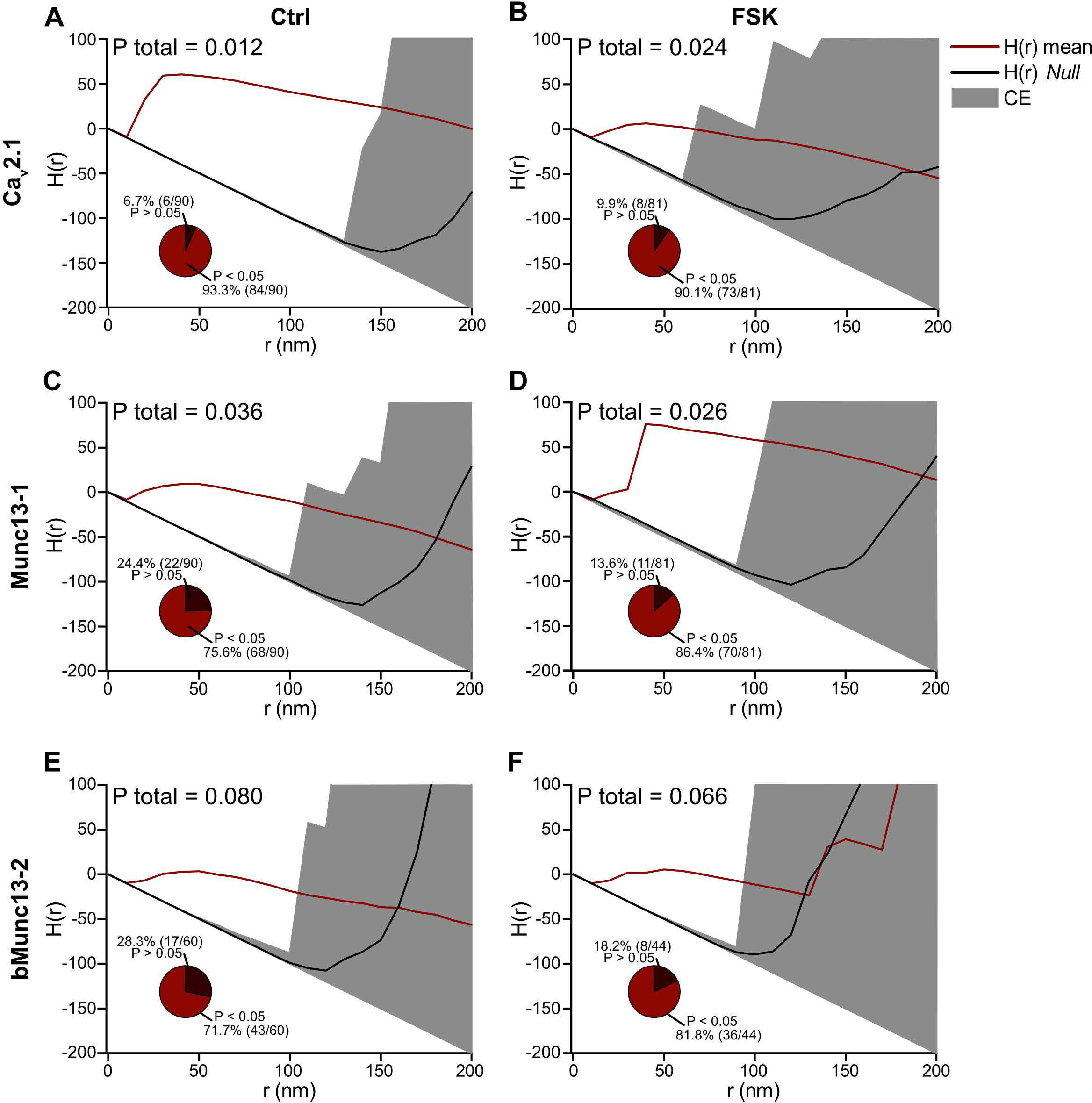
related to Figure 4**. Univariate Ripley-function of Ca_V_2.1s and Munc13s proteins particles in MFBs before and after 5-min forskolin treatment.** **(A** and **B)** Univariate H(r) function of Ca_V_2.1 in DMSO control (A) and after forskolin (B); n AZ = 90 and 81 respectively, total P-value on the figure. Red line indicate population mean, black line random distribution, and gray area - confidence envelopes (CE). Insert: pie chart of statistical significance of MAD test of single AZ. **(C** and **D)** Univariate H(r) function of Munc13-1 in DMSO control (C) and after forskolin (D); n AZ = 90 and 81 respectively, total P-value on the figure. Red line indicates population mean, black line – random distribution, and gray area – confidence envelopes (CE). Insert: pie chart of statistical significance of MAD test of single AZ. **(E** and **F)** Univariate H(r) function of bMunc13-2 in DMSO control (E) and after forskolin (F); n AZ = 60 and 44 respectively, total P-value on the figure. Red line indicates population mean, black line – random distribution, and gray area – confidence envelopes (CE). Insert: pie chart of statistical significance of MAD test of single AZ.

**Figure S6.**
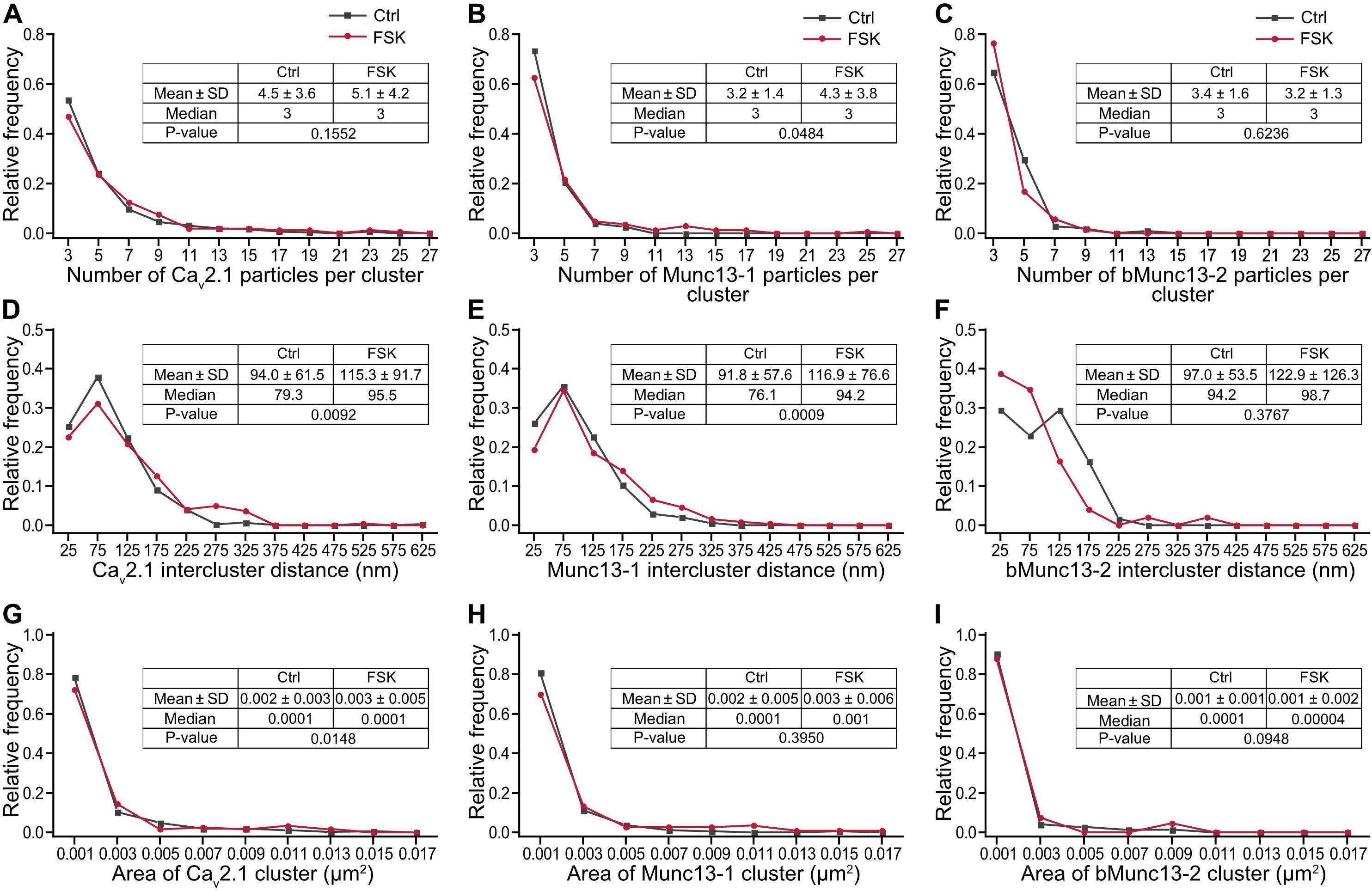
related to Figure 4**. Characteristics of Ca_V_2.1 and Munc13 clusters before and after 5-min FSK application.** **(A–C)** Histogram of relative frequency distribution of number of particles per Ca_V_2.1 (A), Munc13-1 (B), and bMunc13-2 (C) cluster in DMSO control (“Ctrl”, gray) and after forskolin (“FSK”, red). **(D–F)** Histogram of the relative frequency distribution of the minimal distance between Ca_V_2.1 (D), Munc13-1 (E), and bMunc13-2 (F) clusters, color scheme is identical to (A–C). **(G–I)** Histogram of relative frequency distribution of area of each cluster of Ca_V_2.1 (G), Munc13-1 (H), and bMunc13-2 (I) particles, color scheme is identical to (A–C).

**Figure S7.**
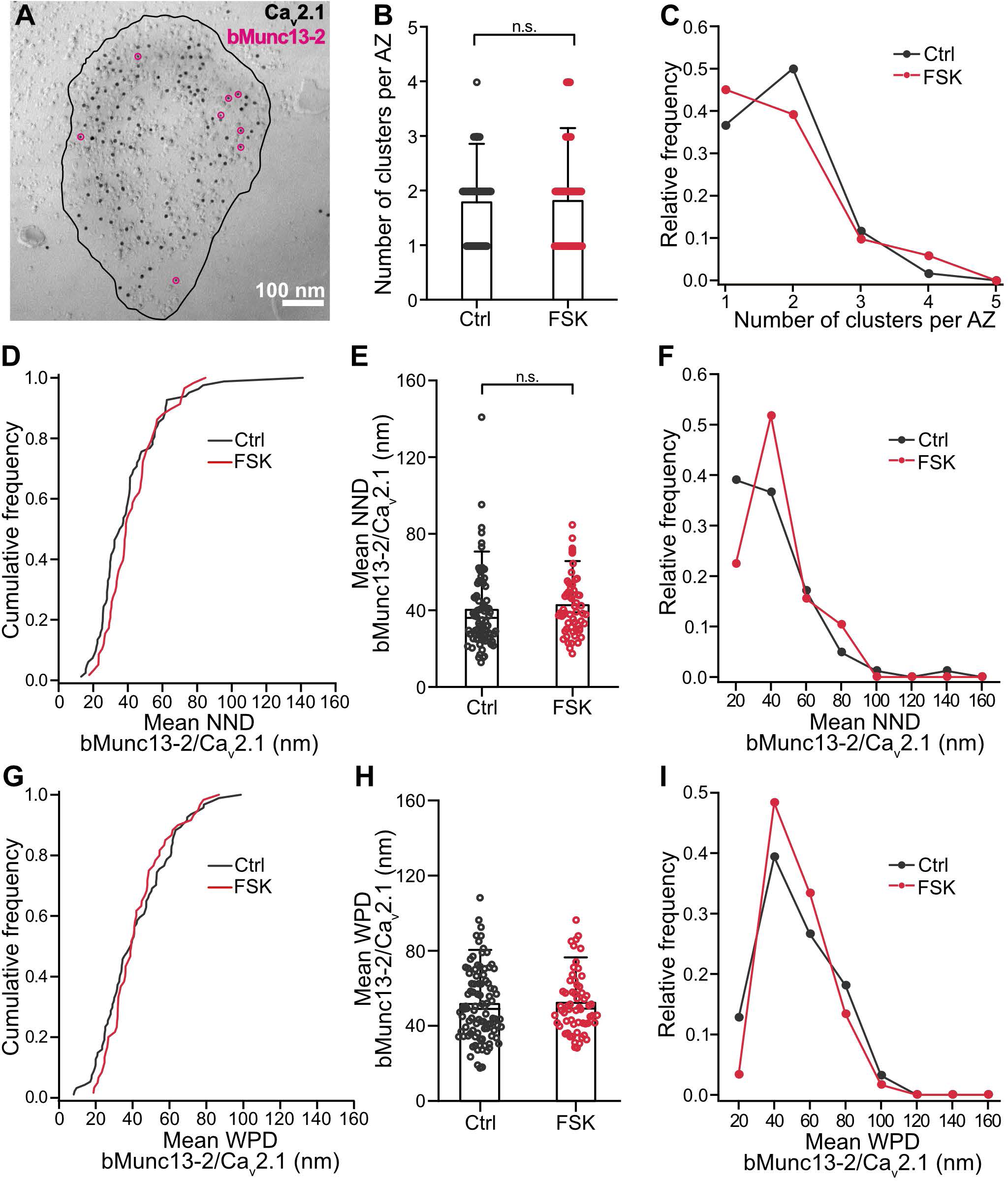
related to Figure 4**. bMunc13-2 shows no alterations in distribution during chemical potentiation.** (A) Example TEM micrograph of freeze-fractured replica of acute slices showing putative MFB AZ (black line) co-labeled against Ca_V_2.1 (black) and bMunc13-2 (pink empty circles). Scale bar size is indicated on the figure. (B) Scatter plot of the number of bMunc13-2 clusters per AZ in DMSO control (“Ctrl”, gray) and after forskolin (“FSK”, red). Bars and whiskers show mean + SD. Horizontal black lines indicate median values. (C) Relative frequency distribution of data displayed in B, color scheme is identical to (B). (D) Cumulative plots of mean NND between experimental bMunc13-2 and Ca_V_2.1 point patterns, color scheme is identical to (B). (E) Scatter plot of the mean NND between bMunc13-2 and Ca_V_2.1 in DMSO control (“Ctrl”, gray) and after FSK treatment (“FSK”, red). Bars and whiskers show mean + SD. Horizontal black lines indicate median values. (F) Relative frequency distribution of data displayed in E, color scheme is identical to (B). (G) Cumulative plots of mean WPDs between experimental bMunc13-2 and Ca_V_2.1 point patterns, color scheme is identical to (B). (H) Summary bar graph of the mean WPDs between bMunc13-2 and Ca_V_2.1, color scheme is identical to (B). Bars and whiskers show mean + SD. Horizontal black lines indicate median values. (I) Relative frequency distribution of data displayed in H, color scheme is identical to (B).

**Table S1.** **related to** **Figure 4. Nonrandom distribution of Ca_v_2.1, Munc13-1 and bMunc13-2 proteins before and after 5-min forskolin treatment**

| | Number of particles per<br>0.1 $\mu\text{m}^2$ AZ | Mean | SD | Median | n AZ<br>(N mice) | p-value |
| --- | --- | --- | --- | --- | --- | --- |
| **Ca<sub>v</sub>2.1** | Control | 36.2 | 12.3 | 35.0 | 130 (5) | 0.3598 |
|  | Forskolin | 38.8 | 15.3 | 38.8 | 52 (3) |  |
| **Munc13-1** | Control | 26.0 | 12.5 | 23.4 | 66 (3) | 0.0689 |
|  | Forskolin | 30.1 | 13.35 | 28.0 | 52 (3) |  |
| **bMunc13-2** | Control | 22.9 | 11.2 | 21.2 | 65 (3) | 0.9543 |
|  | Forskolin | 22.6 | 11.62 | 21.4 | 49 (3) |  |
|  | **Mean NND (nm)** |  |  |  |  |  |
| **Ca<sub>v</sub>2.1** | Control | 26.6 | 6.2 | 25.9 | 106 (5) | < 0.0001 |
|  | Control *Null* | 40.3 | 15.0 | 36.8 | 106 |  |
|  | Forskolin | 25.0 | 4.8 | 23.9 | 52 (3) | < 0.0001 |
|  | Forskolin *Null* | 37.2 | 13.2 | 32.9 | 52 |  |
| **Munc13-1** | Control | 37.5 | 32.3 | 28.7 | 99 (3) | < 0.0001 |
|  | Control *Null* | 59.0 | 33.4 | 50.0 | 99 |  |
|  | Forskolin | 28.9 | 8.1 | 27.7 | 51 (3) | < 0.0001 |
|  | Forskolin *Null* | 37.2 | 13.2 | 32.9 | 52 |  |
| **bMunc13-2** | Control | 41.2 | 23.4 | 36.2 | 92 (3) | < 0.0001 |
|  | Control *Null* | 69.5 | 39.9 | 59.0 | 92 |  |
|  | Forskolin | 33.7 | 12.0 | 31.3 | 48 (3) | < 0.0001 |
|  | Forskolin *Null* | 59.2 | 37.1 | 52.2 | 48 |  |

**Table S2.** **related to** **Figure 4. Spatial distribution of Cav2.1, Munc13-1, and bMunc13-2 proteins after 5-min forskolin treatment**

|  | Number of clusters per AZ | Mean | SD | Median | n AZ (N mice) | p-value |
| --- | --- | --- | --- | --- | --- | --- |
| **Ca<sub>v</sub>2.1** | Control | 2.7 | 1.1 | 3 | 139 (5) | 0.1595 |
|  | Forskolin | 2.8 | 1.4 | 3 | 81 (3) |  |
| **Munc13-1** | Control | 2.4 | 1.1 | 2 | 119 (5) | 0.0088 |
|  | Forskolin | 3.0 | 1.4 | 3 | 78 (3) |  |
| **bMunc13-2** | Control | 1.8 | 0.7 | 2 | 60 (3) | 0.8124 |
|  | Forskolin | 1.8 | 0.9 | 2 | 42 (3) |  |
|  | **Mean NND (nm)** |  |  |  |  |  |
| **Munc13-1+ Ca<sub>v</sub>2.1** | Control | 38.8 | 19.5 | 32.3 | 92 (5) | 0.0275 |
|  | Forskolin | 32.2 | 12.9 | 30.8 | 87(3) |  |
| **bMunc13-2 + Ca<sub>v</sub>2.1** | Control | 40.3 | 20.3 | 36.1 | 94 (3) | 0.1231 |
|  | Forskolin | 42.7 | 15.3 | 38.9 | 59 (3) |  |
|  | **Mean WPD (nm)** |  |  |  |  |  |
| **Munc13-1+ Ca<sub>v</sub>2.1** | Control | 47.3 | 20.6 | 44.4 | 92 (5) | 0.0706 |
|  | Forskolin | 40.6 | 13.9 | 38.7 | 86(3) |  |
| **bMunc13-2 + Ca<sub>v</sub>2.1** | Control | 51.6 | 19.2 | 49.0 | 94 (3) | 0.6687 |
|  | Forskolin | 52.2 | 16.1 | 49.1 | 60 (3) |  |

**Table S3.** **related to Figure5. Spatial distribution of Munc13-1 after treatment with the PKA inhibitor H-89**

|  | Number of clusters per AZ | Mean | SD | Median | n AZ (N mice) |
| --- | --- | --- | --- | --- | --- |
| **Munc13-1** | Control | 2.4 | 1.1 | 2 | 62 (3) |
|  | H-89 | 2.6 | 1.4 | 2 | 52 (3) |
|  | H-89 + Forskolin | 2.4 | 1.3 | 2 | 44 (3) |
|  | **Mean NND (nm)** |  |  |  |  |
| **Munc13-1 + Cav2.1** | Control | 38.8 | 19.5 | 32.3 | 92 (5) |
|  | H-89 | 70.2 | 37.6 | 63.3 | 53 (3) |
|  | H-89 + Forskolin | 43.0 | 26.2 | 37.8 | 42 (3) |
|  | **Mean WPD (nm)** |  |  |  |  |
| **Munc13-1 + Cav2.1** | Control | 47.3 | 20.6 | 44.4 | 92 (5) |
|  | H-89 | 78.4 | 36.9 | 73.0 | 53 (3) |
|  | H-89 + Forskolin | 51.2 | 26.9 | 44.9 | 44(3) |

**Table S4.** **related to Figures 1–4. Comparison of RRP SV size, docked SV pool size, and number of primed SVs at hippocampal MFBs in control and during chemical potentiation**

|  | Ctrl |  | Chemical potentiation |  |
| --- | --- | --- | --- | --- |
|  | Single AZ | Whole MFB | Single AZ | Whole MFB |
| **RRP SV\*** | 1.6 | 47 | 4 | 110 |
| **Docked SVs<sup>♦</sup>** | 13 | 377 | 21 | 609 |
| **Primed SVs<sup>♠</sup>** | 4.2 | 122 | 4.8 | 139 |

\* To determine the number of SV per AZ, which are part of RRP, we divided mean RRP (nA) by mean mEPSCs amplitude (pA). To estimate the number of RRP of SVs in the whole MFB, we multiplied single AZ number by 29 (average number of AZs per P28-old large MFB, Rollenhagen et al., 2007).
♦ To quantify number of docked SVs per AZ, we estimated the area of single profile ( $0.007 \mu\text{m}^2$ ) and extrapolated the number of docked SV per profile to $0.12 \mu\text{m}^2$ (average AZ size in P28-old MFB, Rollenhagen et al., 2007). Next, we multiplied single AZ number by 29 (average number of AZs per postnatal day 28 large MFB, Rollenhagen et al., 2007).
♠ Primed SVs per AZ were assumed to be equal to number of Munc13-1 and bMunc13-2 clusters. Next, we multiplied the single-AZ number by 29 (average number of AZs postnatal day 28 large MFB, Rollenhagen et al., 2007).
\*, ♦, ♠ Numbers of RRP, docked SVs, and primed SVs were determined in mice; the number of AZs average AZ size were determined in rats (Rollenhagen et al., 2007). Thus, effects of species differences on our estimates cannot be excluded.

## REFERENCES

1. Abbott LF, Regehr WG (2004) Synaptic computation. Nature 431, 796–803.

2. Adler EM, Augustine GJ, Duffy SN, Charlton MP (1991) Alien intracellular calcium chelators attenuate neurotransmitter release at the squid giant synapse. J Neurosci 11, 1496–1507.

3. Alabi AA, Tsien RW (2012) Synaptic vesicle pools and dynamics. Cold Spring Harb Perspect Biol 4, a013680.

4. Aldahabi M, Balint F, Holderith N, Lorincz A, Reva M, Nusser Z (2022) Different priming states of synaptic vesicles underlie distinct release probabilities at hippocampal excitatory synapses. Neuron 110, 4144–4161.e7.

5. Augustin I, Betz A, Herrmann C, Jo T, Brose N (1999a) Differential expression of two novel Munc13 proteins in rat brain. Biochem J 337, 363–371.

6. Augustin I, Rosenmund C, Südhof TC, Brose N (1999b) Munc13-1 is essential for fusion competence of glutamatergic synaptic vesicles. Nature 400, 457–461.

7. Banerjee A, Imig C, Balakrishnan K, Kershberg L, Lipstein N, Uronen RL, Wang J, Cai X, Benseler F, Rhee JS, Cooper BH, Liu C, Wojcik SM, Brose N, Kaeser PS (2022) Molecular and functional architecture of striatal dopamine release sites. Neuron 110, 248–265.e9.

8. Betz A, Thakur P, Junge HJ, Ashery U, Rhee JS, Scheuss V, Rosenmund C, Rettig J, Brose N (2001) Functional interaction of the active zone proteins Munc13-1 and RIM1 in synaptic vesicle priming. Neuron 30,183–196.

9. Bischofberger J, Engel D, Frotscher M, Jonas P (2006a) Timing and efficacy of transmitter release at mossy fiber synapses in the hippocampal network. Pflügers Arch 453, 361–372.

10. Bischofberger J, Engel D, Li L, Geiger JRP, Jonas P (2006b) Patch-clamp recording from mossy fiber terminals in hippocampal slices. Nat Protoc 1, 2075–2081.

11. Borges-Merjane C, Kim O, Jonas P (2020) Functional electron microscopy, “Flash and Freeze,” of identified cortical synapses in acute brain slices. Neuron 105, 992–1006.e6.

12. Breustedt J, Gundlfinger A, Varoqueaux F, Reim K, Brose N, Schmitz D (2010) Munc13-2 differentially affects hippocampal synaptic transmission and plasticity. Cereb Cortex 20, 1109– 1120.

13. Brose N, Hofmann K, Hata Y, Südhof TC (1995) Mammalian homologues of Caenorhabditis elegans unc-13 gene define novel family of C2-domain proteins. J Biol Chem 270, 25273– 25280.

14. Bucurenciu I, Kulik A, Schwaller B, Frotscher M, Jonas P (2008) Nanodomain coupling between Ca^2+^ channels and Ca^2+^ sensors promotes fast and efficient transmitter release at a cortical GABAergic synapse. Neuron 57, 536–545.

15. Camacho M, Basu J, Trimbuch T, Chang S, Pulido-Lozano C, Chang SS, Duluvova I, Abo- Rady M, Rizo J, Rosenmund C (2017) Heterodimerization of Munc13 C_2_A domain with RIM regulates synaptic vesicle docking and priming. Nat Commun 8, 15293.

16. Castillo PE (2012) Presynaptic LTP and LTD of excitatory and inhibitory synapses. Cold Spring Harb Perspect Biol 4, a005728.

17. Castillo PE, Weisskopf MG, Nicoll RA (1994) The role of Ca^2+^ channels in hippocampal mossy fiber synaptic transmission and long-term potentiation. Neuron 12, 261–269.

18. Chen JJ, Kaufmann WA, Chen C, Arai I, Kim O, Shigemoto R, Jonas P (2023) Developmental transformation of Ca^2+^ channel–vesicle nanotopography at a central GABAergic synapse. Neuron, accepted for publication.

19. Chen Z, Cooper B, Kalla S, Varoqueaux F, Young SM Jr (2013) The Munc13 proteins differentially regulate readily releasable pool dynamics and calcium-dependent recovery at a central synapse. J Neurosci 33, 8336–8351.

20. Clements JD, Bekkers JM (1997) Detection of spontaneous synaptic events with an optimally scaled template. Biophys J 73, 220–229.

21. Delvendahl I, Vyleta NP, von Gersdorff H, Hallermann S (2016) Fast, temperature-sensitive and clathrin-independent endocytosis at central synapses. Neuron 90, 492–498.

22. Eggermann E, Bucurenciu I, Goswami SP, Jonas P (2012) Nanodomain coupling between Ca²⁺ channels and sensors of exocytosis at fast mammalian synapses. Nat Rev Neurosci 13, 7–21.

23. Emperador-Melero J, Kaeser PS (2020) Assembly of the presynaptic active zone. Curr Opin Neurobiol 63, 95–103.

24. Ester M, Kriegel H, Sander J, Xu X (1996) A density-based algorithm for discovering clusters in large spatial databases with noise. Knowledge Discovery and Data Mining AAAI Press, 226–231.

25. Fernandes HB, Riordan S, Nomura T, Remmers CL, Kraniotis S, Marshall JJ, Kukreja L, Vassar R, Contractor A (2015) Epac2 mediates cAMP-dependent potentiation of neurotransmission in the hippocampus. J Neurosci 35, 6544–6553.

26. Frotscher M, Jonas P, Sloviter RS (2006) Synapses formed by normal and abnormal hippocampal mossy fibers. Cell Tissue Res 326, 361–367.

27. Fukaya R, Hirai H, Sakamoto H, Hashimotodani Y, Hirose K, and Sakaba T (2023) Increased vesicle fusion competence underlies long-term potentiation at hippocampal mossy fiber synapses. Sci Adv 9, eadd3616.

28. Fukaya R, Maglione M, Sigrist SJ, Sakaba T (2021) Rapid Ca^2+^ channel accumulation contributes to cAMP-mediated increase in transmission at hippocampal mossy fiber synapses. Proc Natl Acad Sci USA 118, e2016754118.

29. Grushin K, Kalyana Sundaram RV, Sindelar CV, Rothman JE (2022) Munc13 structural transitions and oligomers that may choreograph successive stages in vesicle priming for neurotransmitter release. Proc Natl Acad Sci USA 119, e2121259119.

30. Hagiwara A, Fukazawa Y, Deguchi-Tawarada M, Ohtsuka T, Shigemoto R (2005) Differential distribution of release-related proteins in the hippocampal CA3 area as revealed by freeze- fracture replica labeling. J Comp Neurol 489, 195–216.

31. Hallermann S, Pawlu C, Jonas P, Heckmann M (2003) A large pool of releasable vesicles in a cortical glutamatergic synapse. Proc Natl Acad Sci USA 100, 8975–8980.

32. Hansen N, Manahan-Vaughan D (2015) Hippocampal long-term potentiation that is elicited by perforant path stimulation or that occurs in conjunction with spatial learning is tightly controlled by beta-adrenoreceptors and the locus coeruleus. Hippocampus 25, 1285–1298.

33. Harris KM (2020) Synaptic odyssey. J Neurosci 40, 61–80.

34. Henze DA, Wittner L, Buzsáki G (2002) Single granule cells reliably discharge targets in the hippocampal CA3 network *in vivo*. Nat Neurosci 5, 790–795.

35. Hilton BJ, Husch A, Schaffran B, Lin TC, Burnside ER, Dupraz S, Schelski M, Kim J, Müller JA, Schoch S, Imig C, Brose N, Bradke F (2022) An active vesicle priming machinery suppresses axon regeneration upon adult CNS injury. Neuron 110, 51–69.e7.

36. Holler S, Köstinger G, Martin KAC, Schuhknecht GFP. Stratford KJ (2021) Structure and function of a neocortical synapse. Nature 591, 111–116.

37. Hopkins WF, Johnston D (1988) Noradrenergic enhancement of long-term potentiation at mossy fiber synapses in the hippocampus. J Neurophysiol 59, 667–687.

38. Huang YY, Kandel ER (1996) Modulation of both the early and the late phase of mossy fiber LTP by the activation of beta-adrenergic receptors. Neuron 16, 611–617.

39. Huang YY, Li XC, Kandel ER (1994) cAMP contributes to mossy fiber LTP by initiating both a covalently mediated early phase and macromolecular synthesis-dependent late phase. Cell 79, 69–79.

40. Imig C, López-Murcia FJ, Maus L, García-Plaza IH, Mortensen LS, Schwark M, Schwarze V, Angibaud J, Nägerl UV, Taschenberger H, Brose N, Cooper BH (2020) Ultrastructural imaging of activity-dependent synaptic membrane-trafficking events in cultured brain slices. Neuron 108, 843–860.e8.

41. Imig C, Min S-W, Krinner S, Arancillo M, Rosenmund C, Südhof TC, Rhee J, Brose N, Cooper BH (2014) The morphological and molecular nature of synaptic vesicle priming at presynaptic active zones. Neuron 84, 416–431.

42. Jackson MB, Redman SJ (2003) Calcium dynamics, buffering, and buffer saturation in the boutons of dentate granule-cell axons in the hilus. J Neurosci 23, 1612–1621.

43. Jonas P, Major G, Sakmann B (1993) Quantal components of unitary EPSCs at the mossy fibre synapse on CA3 pyramidal cells of rat hippocampus. J Physiol 472, 615–663.

44. Josselyn SA, Tonegawa S (2020) Memory engrams: Recalling the past and imagining the future. Science 367, eaaw4325.

45. Kaeser PS, Regehr WG (2017) The readily releasable pool of synaptic vesicles. Curr Opin Neurobiol 43, 63–70.

46. Karlocai MR, Heredi J, Benedek T, Holderith N, Lorincz A, Nusser Z (2021) Variability in the Munc13-1 content of excitatory release sites. eLife 10, e67468.

47. Katz, B (1969) The Release of Neural Transmitter Substances. Liverpool University Press, Liverpool.

48. Kaufman AM, Geiller T, Losonczy A (2020) A role for the locus coeruleus in hippocampal CA1 place cell reorganization during spatial reward learning. Neuron 105, 1018–1026.e4.

49. Kawabe H, Mitkovski M, Kaeser PS, Hirrlinger J, Opazo F, Nestvogel D, Kalla S, Fejtova A, Verrier SE, Bungers SR, Cooper BH, Varoqueaux F, Wang Y, Nehring RB, Gundelfinger ED, Rosenmund C, Rizzoli SO, Südhof TC, Rhee J-S, Brose N (2017) ELKS1 localizes the synaptic vesicle priming protein bMunc13-2 to a specific subset of active zones. J Cell Biol 216, 1143–1161.

50. Kempadoo KA, Mosharov EV, Choi SJ, Sulzer D, Kandel ER (2016) Dopamine release from the locus coeruleus to the dorsal hippocampus promotes spatial learning and memory. Proc Natl Acad Sci USA 113, 14835–14840.

51. Kiskowski MA, Hancock JF, Kenworthy AK (2009) On the use of Ripley’s K-function and its derivatives to analyze domain size. Biophys J 97, 1095–1103.

52. Kobayashi K, Shikano K, Kuroiwa M, Horikawa M, Ito W, Nishi A, Segi-Nishida E, Suzuki H (2022) Noradrenaline activation of hippocampal dopamine D_1_ receptors promotes antidepressant effects. Proc Natl Acad Sci USA 119, e2117903119.

53. Kobayashi K, Suzuki H (2007) Dopamine selectively potentiates hippocampal mossy fiber to CA3 synaptic transmission. Neuropharmacology 52, 552–561.

54. Kobbersmed JRL, Grasskamp AT, Jusyte M, Böhme MA, Ditlevsen S, Sørensen JB, Walter AM (2020) Rapid regulation of vesicle priming explains synaptic facilitation despite heterogeneous vesicle:Ca^2+^ channel distances. eLife 9, e51032.

55. Lashley KS (1950) In search of the engram. In Society for Experimental Biology, Physiological mechanisms in animal behavior. (Society’s Symposium IV.), 454–482. Academic Press.

56. Lawrence JJ, Grinspan ZM, McBain CJ (2004) Quantal transmission at mossy fibre targets in the CA3 region of the rat hippocampus. J Physiol 554, 175–193.

57. Li L, Bischofberger J, Jonas P (2007) Differential gating and recruitment of P/Q-, N-, and R- type Ca^2+^ channels in hippocampal mossy fiber boutons. J Neurosci 27, 13420–13429.

58. Limbach C, Laue MM, Wang X, Hu B, Thiede N, Hultqvist G, Kilimann MW (2011) Molecular in situ topology of Aczonin/Piccolo and associated proteins at the mammalian neurotransmitter release site. Proc Natl Acad Sci USA 108, E392–401.

59. Lipstein N, Chang S, Lin K-H, López-Murcia FJ, Neher E, Taschenberger H, Brose N (2021) Munc13-1 is a Ca^2+^-phospholipid-dependent vesicle priming hub that shapes synaptic short- term plasticity and enables sustained neurotransmission. Neuron 109, 3980–4000.e7.

60. Lipstein N, Sakaba T, Cooper BH, Lin K-H, Strenzke N, Ashery U, Rhee J-S, Taschenberger H, Neher E, Brose N (2013) Dynamic control of synaptic vesicle replenishment and short-term plasticity by Ca^2+^-calmodulin-Munc13-1 signaling. Neuron 79, 82–96.

61. López-García JC, Arancio O, Kandel ER, Baranes D (1996) A presynaptic locus for long-term potentiation of elementary synaptic transmission at mossy fiber synapses in culture. Proc Natl Acad Sci USA 93, 4712–4717.

62. Loy R, Koziell DA, Lindsey JD, Moore RY (1980) Noradrenergic innervation of the adult rat hippocampal formation. J Comp Neurol 189, 699–710.

63. Lu J, Machius M, Dulubova I, Dai H, Südhof TC, Tomchick DR, Rizo J (2006) Structural basis for a Munc13-1 homodimer to Munc13-1/RIM heterodimer switch. PLoS Biol 4, e192.

64. Maccaferri G, Tóth K, McBain CJ (1998) Target-specific expression of presynaptic mossy fiber plasticity. Science 279, 1368–1371.

65. Man KN, Imig C, Walter AM, Pinheiro PS, Stevens DR, Rettig J, Sørensen JB, Cooper BH, Brose N, Wojcik SM (2015) Identification of a Munc13-sensitive step in chromaffin cell large dense-core vesicle exocytosis. Elife 4, e10635.

66. Maus L, Lee C, Altas B, Sertel SM, Weyand K, Rizzoli SO, Rhee J, Brose N, Imig C, Cooper BH (2020) Ultrastructural correlates of presynaptic functional heterogeneity in hippocampal synapses. Cell Rep 30, 3632–3643.e8.

67. Monday HR, Younts TJ, Castillo PE (2018) Long-term plasticity of neurotransmitter release: Emerging mechanisms and contributions to brain function and disease. Annu Rev Neurosci 41, 299–322.

68. Moudy AM, Kunkel DD, Schwartzkroin P (1993) Development of dopamine-beta- hydroxylase- positive fiber innervation of the rat hippocampus. Synapse 15, 307–318.

69. Midorikawa M, Sakaba T (2017) Kinetics of releasable synaptic vesicles and their plastic changes at hippocampal mossy fiber synapses. Neuron 96, 1033–1040.e3.

70. Müller JA, Betzin J, Santos-Tejedor J, Mayer A, Oprişoreanu AM, Engholm-Keller K, Paulußen I, Gulakova P, McGovern TD, Gschossman LJ, Schönhense E, Wark JR, Lamprecht A, Becker AJ, Waardenberg AJ, Graham ME, Dietrich D, Schoch S (2022) A presynaptic phosphosignaling hub for lasting homeostatic plasticity. Cell Rep 39, 110696.

71. Nagy G, Reim K, Matti U, Brose N, Binz T, Rettig J, Neher E, Sørensen JB (2004) Regulation of releasable vesicle pool sizes by protein kinase A-dependent phosphorylation of SNAP-25. Neuron 41(3), 417–429

72. Neher E (2015) Merits and limitations of vesicle pool models in view of heterogeneous populations of synaptic vesicles. Neuron 87, 1131–1142.

73. Neher E (2017) Some subtle lessons from the calyx of Held synapse. Biophys J 112, 215– 223.

74. Neher E, Brose N (2018) Dynamically primed synaptic vesicle states: Key to understand synaptic short-term plasticity. Neuron 100, 1283–1291.

75. Nicoll RA, Schmitz D (2005) Synaptic plasticity at hippocampal mossy fibre synapses. Nat Rev Neurosci 6, 863–876.

76. O’Dell TJ, Connor SA, Guglietta R, Nguyen PV (2015) β-Adrenergic receptor signaling and modulation of long-term potentiation in the mammalian hippocampus. Learn Mem 22, 461– 471.

77. Orlando M, Dvorzhak A, Bruentgens F, Maglione M, Rost BR, Sigrist SJ, Breustedt J, Schmitz D (2021) Recruitment of release sites underlies chemical presynaptic potentiation at hippocampal mossy fiber boutons. PLoS Biol 19, e3001149.

78. Patzke C, Brockmann MM, Dai J, Gan KJ, Grauel MK, Fenske P, Liu Y, Acuna C, Rosenmund C, Südhof TC (2019) Neuromodulator signaling bidirectionally controls vesicle numbers in human synapses. Cell 179, 498–513.e22.

79. Pauli M, Paul MM, Proppert S, Mrestani A, Sharifi M, Repp F, Kürzinger L, Kollmannsberger P, Sauer M, Heckmann M, Sirén AL (2021) Targeted volumetric single-molecule localization microscopy of defined presynaptic structures in brain sections. Nat Commun Biol 4, 407.

80. Rebola N, Reva M, Kirizs T, Szoboszlay M, Lőrincz A, Moneron G, Nusser Z, DiGregorio DA (2019) Distinct nanoscale calcium channel and synaptic vesicle topographies contribute to the diversity of synaptic function. Neuron 104, 693–710.e9.

81. Rey S, Marra V, Smith C, Staras K (2020) Nanoscale remodeling of functional synaptic vesicle pools in Hebbian plasticity. Cell Rep 30, 2006–2017.e3.

82. Ripley BD (1977) Modelling spatial patterns. J. R. Stat. Soc. B 39, 172–192.

83. Rizzoli SO, Betz WJ (2005) Synaptic vesicle pools. Nat Rev Neurosci 6, 57–69.

84. Rollenhagen A, Sätzler K, Rodríguez EP, Jonas P, Frotscher M, Lübke JHR (2007) Structural determinants of transmission at large hippocampal mossy fiber synapses. J Neurosci 27, 10434–10444.

85. Sakaguchi G, Orita S, Naito A, Maeda M, Igarashi H, Sasaki T, Takai Y. (1998) A novel brain- specific isoform of beta spectrin: isolation and its interaction with Munc13 Biochem Biophys Res Commun 248, 846–851.

86. Sakamoto H, Ariyoshi T, Kimpara N, Sugao K, Taiko I, Takikawa K, Asanuma D, Namiki S, Hirose K (2018) Synaptic weight set by Munc13-1 supramolecular assemblies. Nat Neurosci 21, 41–49.

87. Salin PA, Scanziani M, Malenka RC, Nicoll RA (1996) Distinct short-term plasticity at two excitatory synapses in the hippocampus. Proc Natl Acad Sci USA 93, 13304–13309.

88. Sara SJ (2009) The locus coeruleus and noradrenergic modulation of cognition. Nat Rev Neurosci 10, 211–223.

89. Schindelin J, Arganda-Carreras I, Frise E, Kaynig V, Longair M, Pietzsch T, Preibisch S, Rueden C, Saalfeld S, Schmid B, Tinevez JY, White DJ, Hartenstein V, Eliceiri K, Tomancak P, Cardona A (2012) Fiji: an open-source platform for biological-image analysis. Nat Methods 9, 676–682.

90. Schneggenburger R, Meyer AC, Neher E (1999) Released fraction and total size of a pool of immediately available transmitter quanta at a calyx synapse. Neuron 23, 399–409.

91. Shahoha M, Cohen R, Ben-Simon Y, Ashery U (2022) cAMP-dependent synaptic plasticity at the hippocampal mossy fiber terminal. Front Synaptic Neurosci 14, 861215.

92. Silver RA (2010) Neuronal arithmetic. Nat Rev Neurosci 11, 474–489.

93. Studer D, Zhao S, Chai X, Jonas P, Graber W, Nestel S, Frotscher M (2014) Capture of activity-induced ultrastructural changes at synapses by high-pressure freezing of brain tissue. Nat Protoc 9, 1480–1495.

94. Takeuchi T, Duszkiewicz AJ, Sonneborn A, Spooner PA, Yamasaki M, Watanabe M, Smith CC, Fernández G, Deisseroth K, Greene RW, Morris RGM (2016) Locus coeruleus and dopaminergic consolidation of everyday memory. Nature 537, 357–362.

95. Tan C, de Nola G, Qiao C, Imig C, Born RT, Brose N, Kaeser PS (2022) Munc13 supports fusogenicity of non-docked vesicles at synapses with disrupted active zones. Elife 11, e79077.

96. Tong G, Malenka RC, Nicoll RA (1996) Long-term potentiation in cultures of single hippocampal granule cells: a presynaptic form of plasticity. Neuron 16, 1147–1157.

97. Vandael D, Borges-Merjane C, Zhang X, Jonas P (2020) Short-term plasticity at hippocampal mossy fiber synapses is induced by natural activity patterns and associated with vesicle pool engram formation. Neuron 107, 509–521.e7.

98. Vandael D, Okamoto Y, Borges-Merjane C, Vargas-Barroso V, Suter BA, Jonas P (2021) Subcellular patch-clamp techniques for single-bouton stimulation and simultaneous pre- and postsynaptic recording at cortical synapses. Nat Protoc 16, 2947–2967.

99. Varoqueaux F, Sigler A, Rhee J-S, Brose N, Enk C, Reim K, Rosenmund C. (2002) Total arrest of spontaneous and evoked synaptic transmission but normal synaptogenesis in the absence of Munc13-mediated vesicle priming. Proc Natl Acad Sci USA 99, 9037–9042.

100. Varoqueaux F, Sons MS, Plomp JJ, Brose N (2005) Aberrant morphology and residual transmitter release at the Munc13-deficient mouse neuromuscular synapse. Mol Cell Biol 25, 5973–5984.

101. Vogt KE, Regehr WG (2001) Cholinergic modulation of excitatory synaptic transmission in the CA3 area of the hippocampus. J Neurosci 21, 75–83.

102. Vyleta NP, Jonas P (2014) Loose coupling between Ca^2+^ channels and release sensors at a plastic hippocampal synapse. Science 343, 665–670.

103. Vyleta NP, Borges-Merjane C, Jonas P (2016) Plasticity-dependent, full detonation at hippocampal mossy fiber-CA3 pyramidal neuron synapses. eLife 5, e17977.

104. Wagatsuma A, Okuyama T, Sun C, Smith LM, Abe K, Tonegawa S (2018) Locus coeruleus input to hippocampal CA3 drives single-trial learning of a novel context. Proc Natl Acad Sci USA 115, E310–E316.

105. Wang SSH, Held RG, Wong MY, Liu C, Karakhanyan A, Kaeser PS (2016) Fusion competent synaptic vesicles persist upon active zone disruption and loss of vesicle docking. Neuron 91, 777–791.

106. Weisskopf MG, Castillo PE, Zalutsky RA, Nicoll RA (1994) Mediation of hippocampal mossy fiber long-term potentiation by cyclic AMP. Science 265, 1878–1882.

107. Wilke SA, Antonios JK, Bushong EA, Badkoobehi A, Malek E, Hwang M, Terada M, Ellisman MH, Ghosh A (2013) Deconstructing complexity: serial block-face electron microscopic analysis of the hippocampal mossy fiber synapse. J Neurosci 33, 507–522.

108. Zarebidaki F, Camacho M, Brockmann MM, Trimbuch T, Herman MA, Rosenmund C (2020) Disentangling the roles of RIM and Munc13 in synaptic vesicle localization and neurotransmission. J Neurosci 40, 9372–9385.

109. Zhang X, Schlögl A, Jonas P (2020) Selective routing of spatial information flow from input to output in hippocampal granule cells. Neuron 107, 1212–1225.e7

